# Evolutionary adaptation from hydrolytic to oxygenolytic catalysis

**DOI:** 10.1101/2023.05.05.539528

**Authors:** Soi Bui, Sara Gil-Guerrero, Peter van der Linden, Philippe Carpentier, Matteo Ceccarelli, Pablo G. Jambrina, Roberto A. Steiner

## Abstract

Protein fold adaptation to novel enzymatic reactions is a fundamental evolutionary process. Cofactor-independent oxygenases degrading *N*-heteroaromatic substrates belong to the α/β-hydrolase (ABH) fold superfamily that typically does not catalyze oxygenation reactions. Here, we have integrated crystallographic analyses at normoxic and hyperoxic conditions with molecular dynamics and quantum mechanical calculations to investigate its prototypic 1-*H*-3-hydroxy-4-oxoquinaldine 2,4-dioxygenase (HOD) member. O_2_ localization to the “oxyanion hole”, where catalysis occurs, is an unfavorable event and the direct competition between dioxygen and water for this site is modulated by the “nucleophilic elbow” residue. A hydrophobic pocket that overlaps with the organic substrate binding site can act as a proximal dioxygen reservoir. Freeze-trap pressurization allowed to determine the structure of the ternary complex with a substrate analogue and O_2_ bound at the oxyanion hole. Theoretical calculations reveal that O_2_ orientation is coupled to the charge of the bound organic ligand. When 1-*H*-3-hydroxy-4-oxoquinaldine is uncharged, O_2_ binds with its molecular axis along the ligand’s C2-C4 direction in full agreement with the crystal structure. Substrate activation triggered by deprotonation of its 3-OH group by the His-Asp dyad, rotates O_2_ by approximately 60 degrees. This geometry maximizes the charge-transfer between the substrate and O_2_ thus weakening the double bond of the latter. Electron density transfer to the O_2_(π*) orbital promotes the formation of the peroxide intermediate via intersystem crossing that is rate-determining. Our work provides a detailed picture of how evolution has repurposed the ABH-fold architecture and its simple catalytic machinery to accomplish metal-independent oxygenation.

**Significance:** Many of the current O_2_-dependent enzymes have evolved from classes that existed prior to the switch from a reducing to an oxidative atmosphere and whose original functions are unrelated to dioxygen chemistry. A group of bacterial dioxygenases belong to the α/β-hydrolase (ABH) fold superfamily that typically does not catalyze oxygenation reactions. These enzymes degrade their *N*-heteroaromatic substrates in a cofactor-independent manner relying only on the simple nucleophile-histidine-acid ABH-fold catalytic toolbox. In this work we show how O_2_ localizes at the catalytic site by taking advantage of multiple strategies that minimize the strong competition by water, the co-substrate in the ancestral hydrolytic enzyme. We also show that substrate activation by the His-Asp catalytic dyad leads a ligand-O_2_ complex that maximizes the electron transfer from the organic substrate to O_2_, thus promoting intersystem crossing and circumventing the spin-forbiddeness of the reaction. Overall, our work explains how evolution has repurposed the ABH-fold architecture and its simple catalytic machinery to accomplish spin-restricted metal-independent oxygenation.

## Introduction

The switch from a reducing to an oxidative atmosphere during our planet’s history spawned the emergence of many O_2_-dependent enzymes, many of which evolved from pre-existing classes, whose original functions are unrelated to dioxygen chemistry.^1^ The α/β-hydrolase (ABH) fold is a common and versatile protein architecture found in all three domains of life. The comprehensive ESTHER (ESTerase, α/β-Hydrolase Enzymes and Relatives) database lists more than 70,000 family members that typically share little sequence similarity but display a common fold and catalytic machinery.^2^ Their core structure is an eight-stranded β-sheet surrounded by α-helices often decorated by additional structural elements, generally referred to as cap or lid domains, that confer functional variation (Figure S1).^3, 4^ Mechanistically, most ABH-fold members rely on the conserved nucleophile-histidine-acidic residue proton-relay system of serine hydrolases. In keeping with this, the majority of ABHs are lipases, proteases, esterases, thioesterases, dehalogenases, and epoxide hydrolases that use H_2_O as co-substrate. However, this is not universal and other members include haloperoxidases, lyases, and even oxygenases that employ H_2_O_2_, HCN, or O_2_, respectively, for catalysis.^5^ This underscores the functional malleability of the ABH fold and its simple catalytic triad.

*Arthrobacter nitroguajacolicus* Rü61a 1-*H*-3-hydroxy-4-oxoquinaldine 2,4-dioxygenase (HOD) and *Pseudomonas putida* 33/1 1-*H*-3-hydroxy-4-oxoquinoline 2,4-dioxygenase (QDO) were the first dioxygenases discovered to belong to the ABH-fold family.^6, 7^ They catalyze the O_2_-dependent breakdown of *N*-heteroaromatic quinolone-type substrates with concomitant carbon monoxide production (Figure 1A). Recently, AqdC1 and AqdC2 from *R. erythropolis*, and AqdC from *M. abscessus* subsp*. abscessus*, have been also identified as ABH-fold dioxygenases that catalyze the same reaction,^8, 9^ and the family of quinolone-type degrading ABH-fold dioxygenases has been significantly expanded by bioinformatic analyses with the identification of more than 150 *bona fide* new members, nine of which have been validated experimentally.^10^ Atypical for oxygenases, these enzymes promote the activation of the triplet ground-state O_2_ molecule without the aid of metal centers or external organic co-factors.^11^ Kinetic measurements indicate that their reaction mechanism relies on a fast initial step, during which the substrates’ 3-hydroxyl group is deprotonated by the His-Asp subset of the triad thus promoting its activation for molecular oxygen attack (step **1**, Figure 1B), whilst the nucleophile is not essential.^12-14^ Consistent with this, in various organisms, a serine to alanine replacement is observed for the ‘nucleophile’ residue (Figure S2). The role of the His-Asp dyad as general base is supported by the crystal structures of HOD and AqdC that have been solved in mechanistically-relevant states.^14, 15^ The dispensability of the nucleophile is in stark contrast with what observed for most ABHs for which the nucleophile is required for the formation of the mandatory covalent acyl-enzyme intermediate.

**Figure 1.**
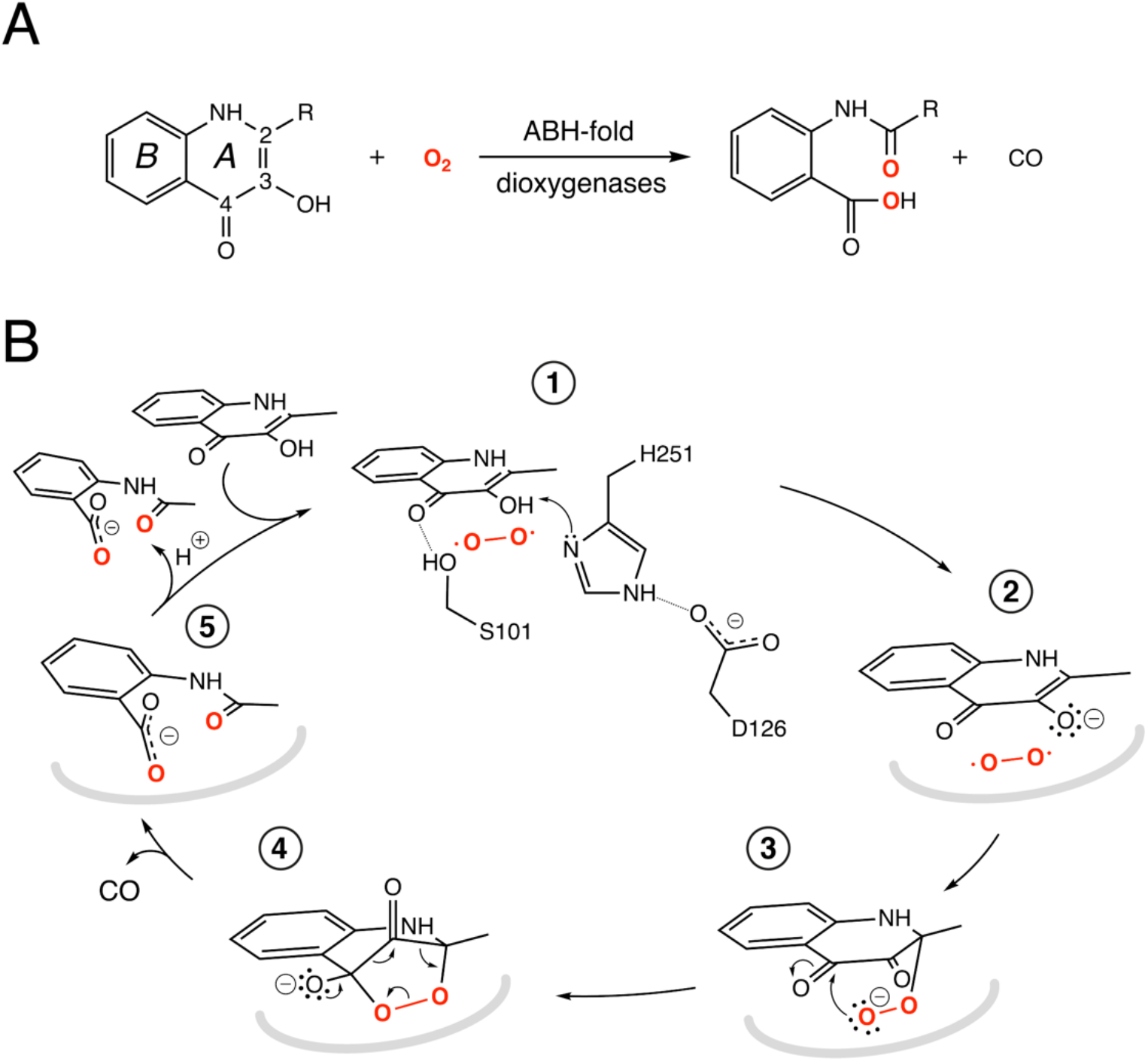
ABH-fold cofactor-indepdendent dioxygenases reaction and mechanism. (A) Scheme of the reaction catalyzed by the bacterial cofactor-independent ABH-fold dioxygenases. HOD, catalyzes primarily conversion of 1-*H*-3-hydroxy-4-oxoquinaldine (R = CH_3_, QND) to *N*-acetylanthranilate (NAA). In the reaction, the *A* heterocycle is disrupted with formation of carbon monoxide as by-product. The compound 2-methyl-quinolin-4(1*H*)-one (MQO) used in this work features -H instead of -OH at position 3; (B) Reaction mechanism as exemplified by HOD. The structure of the anaerobic HOD-QND complex revealed that the 3-OH group of QND is deprotonated by the H251-D126 pair of the ‘nucleophile-histidine-acidic’ ABH-fold triad (steps **1** and **2**). S101 at the sharp structural turn known as the ‘nucleophilic elbow’ further stabilizes the substrate in the active site. In some ABH-fold dioxygenases an alanine residue replaces the serine at the ‘nucleophilic elbow’. The reaction is then assumed to proceed via the peroxide intermediates (**3** and **4**) to the formation of the product NAA (**5**). Hydrogen bonds are represented dotted lines. In reaction steps (**2**-**5**) the protein environment is shown in a simplified manner as a grey curved line.

Cofactor-independent ABH-fold oxygenases are intriguing from a mechanistic viewpoint as the spin-forbidden oxygenation reaction is enabled by a minimalistic catalytic toolbox. Theoretical investigations have put forward different hypotheses for the mechanism that allows to overcome the quantum chemical hurdle of the direct reaction between the singlet-state substrate anion and the triplet-state O_2_.^16, 17^ According to one QM/MM study the rate limiting step of the reaction is the addition of O_2_ to C2 of QND leading to the formation of the peroxide (steps **2**-**3,** Figure 1B) that has been suggested to occur on the triplet-state potential energy surface with a 17 kcal/mol barrier, followed by an intersystem crossing leading to a singlet state.^16^ Another work instead suggested that the triplet-state peroxide intermediate is unlikely to play a role in the mechanism and that catalysis could proceed via a direct electron transfer from the QND anion to O_2,_ followed by radical recombination yielding the peroxide intermediate.^17^ Once the peroxide **3** is formed, subsequent ring closure (step **4**) followed by release of CO and formation of the product (step **5**) are uncontroversial.

The most elusive and interesting aspects of cofactor-independent oxygenation at the ABH-fold are those that involve molecular oxygen for which no direct structural information is currently available. In this work, we have addressed this gap using an integrated computational and experimental framework. This approach has allowed to obtain a complete understanding of the reaction mechanism providing a rationale for how this versatile protein architecture and its simple catalytic machinery tuned for hydrolytic reactions has been successfully redeployed during evolution to carry out O_2_-dependent catalysis.

## Results and discussion

### HOD displays an O_2_ pocket that partly overlaps with that of the organic substrate

To identify possible dioxygen pockets within HOD we initially performed molecular dynamics (MD) simulations. Two independent replica simulations, each 1 μs-long, were carried out for water-solvated HOD in the presence of dioxygen (ten O_2_ molecules in a box of approximately 74×74×74 Å^3^). Identical simulations were also performed without explicit dioxygen. Hereafter, we will refer to simulations with and without explicit O_2_ as OXY-MD and WAT-MD, respectively. In all runs, we observed only minor deviations from the starting crystallographic model (PDB code 2WM2) as expected from the stable ABH fold (Figure S3). In OXY-MD runs, we sampled O_2_ positions every 50 ps mapping the locations most frequently visited sites on this time scale (Figure 2A). These tend to be confined to the core domain and are typically transient pockets afforded by sidechain movements. Amongst the top sites there is however also a portion of the active site (*B*-site in Figure 2A) that is substantially more accessible. This location overlaps with the binding site of the natural substrate, specifically of its hydrophobic *B*-ring portion (Figure 1A). We also compared root-mean-square fluctuations (r.m.s.f.) of the Cα coordinates from the average structures in OXY-MD and WAT-MD runs (Figure S4), a measure of the structural changes during the simulations. It suggests that O_2_ tends to reduce the mobility of the β5-αC ‘nucleophilic elbow’ centered at S101, the portion of the cap domain overhanging it, and the β8-αF’ C-terminal loop hosting the catalytic H251 residue. These regions (highlighted in yellow in Figure 2A) define the interface between the core and cap domains where the active site is located. The basin-shaped portion of the active site that is expected to host O_2_ for its attack on the activated substrate (Figure 2B) is not amongst the most frequented locations on the 50-ps time scale. This O_2_ ‘reactive-site’ (*R-*site) was identified in the crystal structure of the anaerobic HOD-QND complex as a ∼15-Å^3^ cavity opening in front of S101 underneath the substrate’s *A*-ring^14^ and corresponds topologically to the ’oxyanion hole’ that stabilizes the negatively charged tetrahedral transition state in hydrolytic reactions.^4, 18^

**Figure 2.**
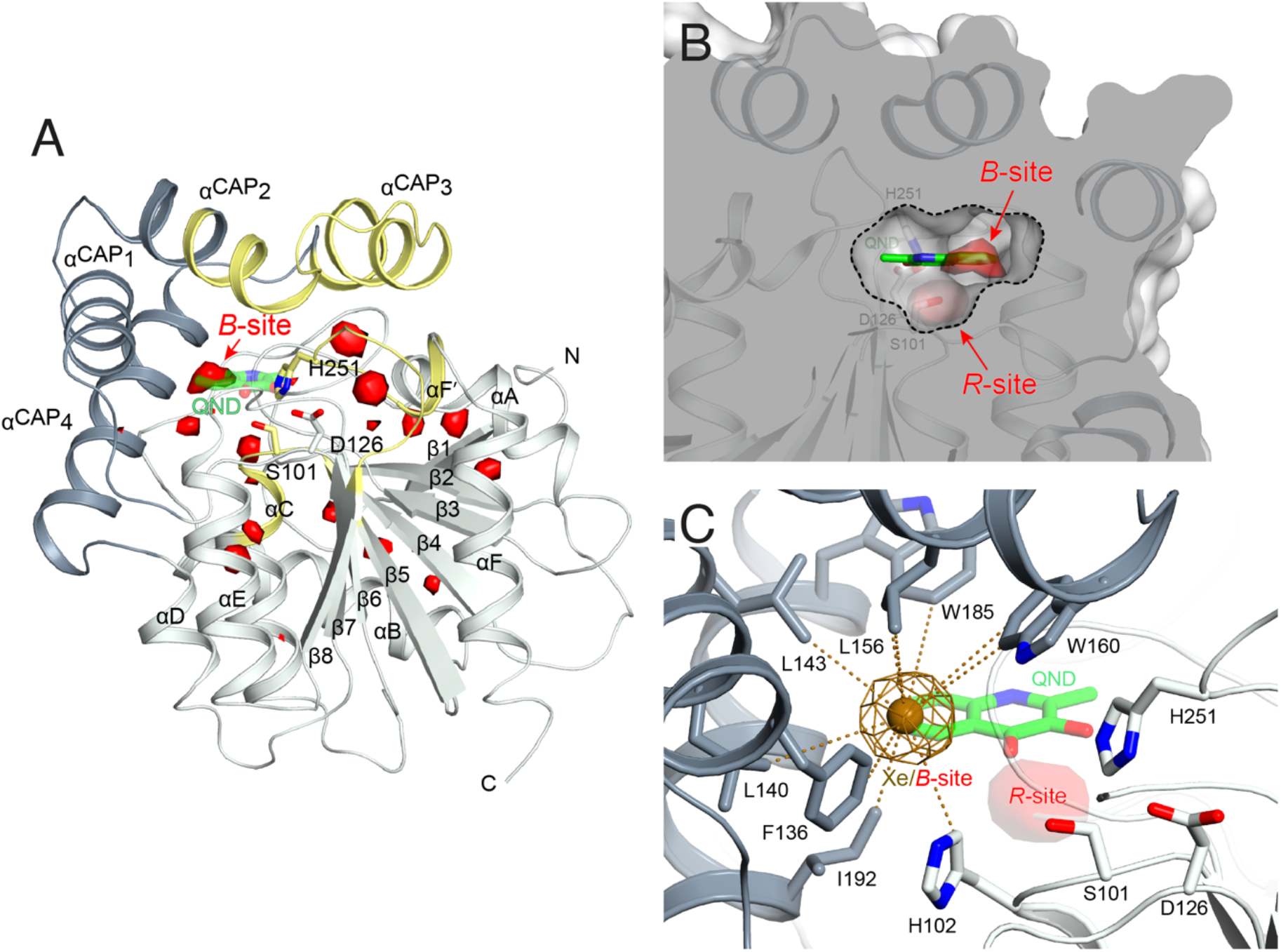
Dioxygen binding sites mapped by MD simulations and xenon pressurization. (A) Cartoon representation of HOD (core and cap domains in light and dark grey, respectively) with high-probability O_2_-sites (isosurface representation in red) as revealed by MD simulations. In yellow are highlighted structural regions whose dynamics is reduced by dioxygen. Most O_2_-sites are temporary pockets generated by protein dynamics within the core domain. One site (*B*-site) is much more accessible and overlaps with the location occupied by the *B*-ring of the QND substrate in the E-S complex. The latter is shown for reference as stick representation with carbon, nitrogen and oxygen atoms colored green, blue and red, respectively. Residues of the catalytic triad are also shown as sticks; (B) Sliced-surface back-view (roughly rotated by 180 degrees around the vertical axis compared to the view in A) showing a cut-through of the active site cavity in the E-S complex (PDB code 2WJ4) with its largest section highlighted by a dotted line). The *B*-site lies within the flat horizontal portion of the active site that hosts the QND substrate. Underneath the substrate’s A-ring heterocycle, the active site is shaped into a basin. The *R*-site (*R* for reactive) that measures approximately 15 Å^3^, positioned in front of the S101 sidechain, is expected to host O_2_ during catalysis; (C) Active site of HOD following xenon pressurization. A Xe atom, shown as a gold sphere with its 2*mF*_o_-*DF*_c_ map at the +1.0σ level as chicken-wire representation, localizes at the *B*-site. Stabilizing sidechains are shown as sticks with atoms within 5.0 Å of Xe highlighted by dotted lines. Residues of the catalytic triad are also shown. All residues are colored according to the domain to which they belong. The bound QND substrate and the *R*-site underneath it are shown for reference.

We next employed xenon pressurization in the crystal state to probe O_2_ sites experimentally. Electron-rich Xe is quite easily detected by crystallographic methods and has been successfully used to visualize hydrophobic O_2_ binding sites in proteins.^19-21^ Pressurization of HOD crystals in a quartz capillary at room-temperature under constant 30 bar Xe atmosphere allowed us to measure X-ray data at 2.9 Å using synchrotron radiation (Table S1). Fourier difference density maps revealed strong peaks consistent with xenon binding at the *B*-site (as it overlaps with the substrate’s *B*-ring in in the HOD-QND complex) of all four HOD molecules present in the asymmetric unit (Figure 2C). Modelling these peaks as water molecules resulted in significant positive residual density post-refinement supporting the notion that Xe occupies this site (Figure S5A). Anomalous maps are also consistent with this assignment (Figure S5B). Occupancy refinement of Xe atoms gives values in the 40-60% range. Xe is stabilized at the *B*-site by the hydrophobic environment of the H102, F136, L140, L143, L156, W160, W185, I192 side chains with the closest atoms at 3.8-5.0 Å from the gas atom (Figure 2C). Although we performed our experiment at room temperature, which compared to cryo-conditions, allows for enhanced protein’s mobility we have not identified additional Xe binding sites in our maps.

Overall, experiments and simulations agree that the ABH-fold HOD dioxygenase features an O_2_ pocket (*B*-site) within the active site at a location that overlaps with the most hydrophobic portion its aromatic substrate.

### S101 at the nucleophilic elbow modulates O_2_/H_2_O stability at the *R*-site

HOD’s active site shape suggests that O_2_ must locate at the *R*-site to react with the bound substrate (Figure 2B). Differently from the hydrophobic *B*-site, the *R*-site is polar and crystallographic studies have shown that a water molecule is often bound at this position with variable occupancy.^13, 14^ In the presence of NaCl at high concentration the *R*-site also stabilizes a chloride ion.^14^ Halide and dioxygen binding sites have been shown to be shared in other O_2_-dependent enzymes.^22-25^

As dioxygen must be in strong competition with H_2_O for the *R*-site and considering the proximity of S101, we wondered whether this residue might play a role in modulating O_2_/H_2_O preference for this location. Serine residues are characterized by three main rotational conformers (rotamers) for their (N-CA-CB-OG1) χ1 torsion angle defined as *plus* (χ1 = +60°), *trans* (χ1 = 180°), and *minus* (χ1 = -60°). Statistical analysis shows that *trans* and *plus* are the least and most frequently observed, respectively, particularly when this residue is not part of either α-helices or β-strands.^26^ We analyzed S101 rotamer preferences in our 1 μs-long substrate-free WAT-MD and OXY-MD simulations and found that O_2_ shifts the distribution from bimodal, in which both *plus* (S101*^p^*) and *trans* (S101*^t^*) rotamers are sampled with equal frequency (Figure 3A), to unimodal in which S101*^t^* dominates (Figure 3B). The *minus* rotamer is never frequently populated. Structurally, the transition from S101*^p^* to S101*^t^*causes the CB-OG bond to reorient itself, switching from a direction pointing directly toward the *R*-site to one ‘running’ tangentially to it following the backbone (Figure 3 inset).

**Figure 3.**
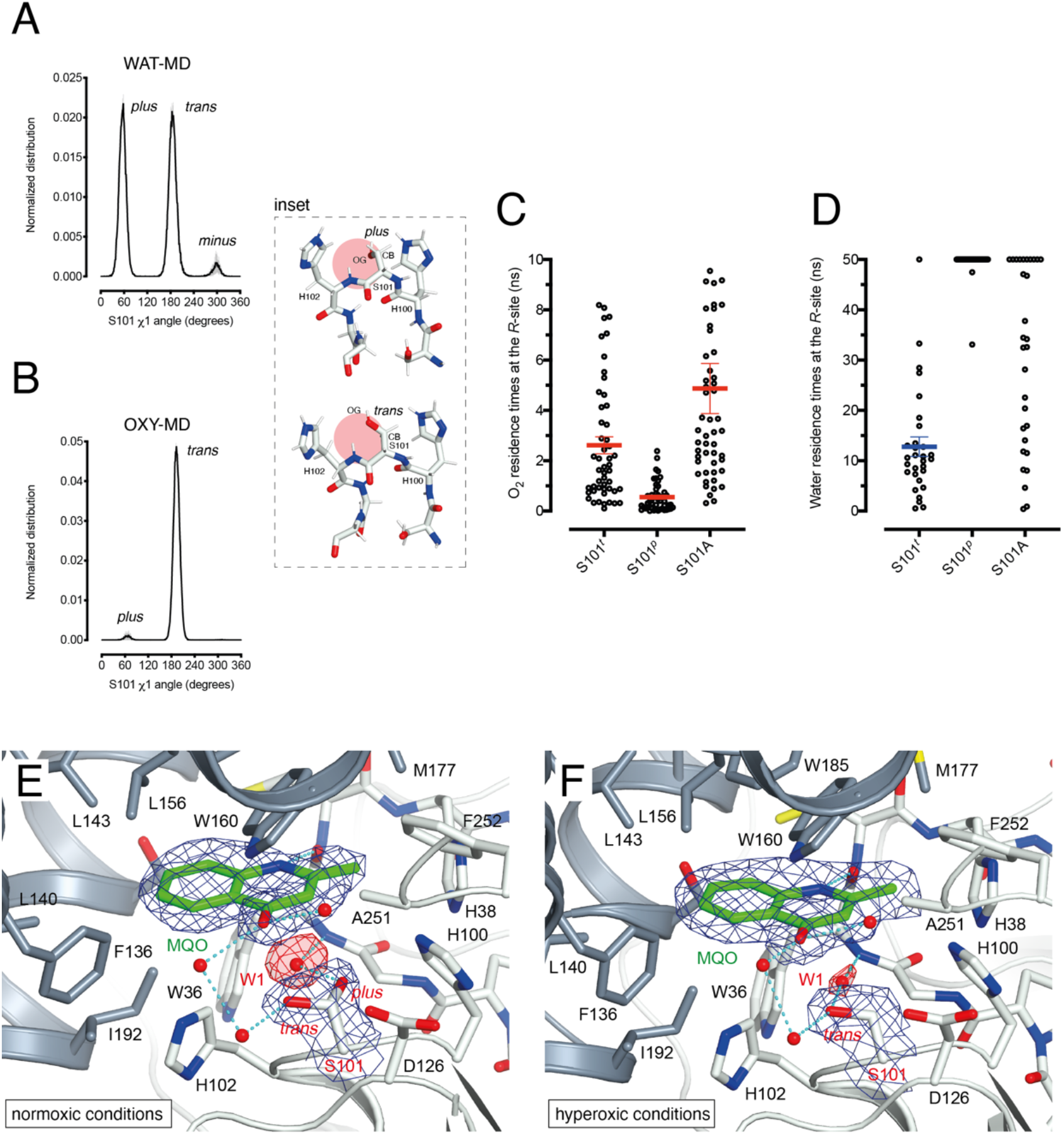
S101 as O_2_/H_2_O modulator at the *R*-site. (A) Normalized distribution of χ1 (N-CA-CB-OG) angles for S101 in MD simulations carried out for HOD without explicit dioxygen. S101 rotamers are defined as: *plus* (χ1 = +60°), *trans* (χ1 = 180°), and *minus* (χ1 = -60°= 300°). Values are the average of two independent MD replicas each 1 μs-long sampled every 50 ps. Standard deviation is shown as light grey bars; (B) Same as A in the presence of explicit dioxygen. The inset provides snapshots during the simulations with the *R*-site represented by the red circle. The CB-OG bond lines the *R*-site for the *trans* rotamer whilst it roughly points into it for the *plus* rotamer; (C) Distribution of residence times for an O_2_ molecule at the *R*-site of the HOD-QND complex with S101 restrained to the *trans* (S101*^t^*), *plus* (S101*^p^*) rotamers or with S101 replaced by an alanine (S101A). A total of 50 independent MD simulations were performed for each system. Mean and s.e.m. values are highlighted; (D) Like C but for a water molecule at the *R*-site. A total of 30 independent MD simulations were performed for each system with a cutoff time of 50 ns. For S101*^t^*, S101*^p^*, and S101A water was observed leaving the *R*-site within the cutoff (29/30), (2/30), (20/30) times, respectively. Mean and s.e.m. values for S101*^t^* are highlighted; (E-F) Active site of the HOD^H251A^-MQO complex under normoxic (E) or hyperoxic (F) conditions. Electron density maps (2*mF*_o_-*DF*_c_) are shown at the +1.0σ level as chicken-wire representation for MQO, the water molecule at the *R*-site (W1) and S101. The latter displays a mixture of *plus* (40%)*/trans* (60%) rotamers under normoxic conditions whilst only the *trans* rotamer is observed under hyperoxic conditions. Electron density for W1 is displayed in red for clarity. H-bond network involving MQO, W1 and S101 is shown as cyan dotted lines.

Next, we turned our attention to the enzyme-substrate (E-S) complex. As expected, a 1 μs-long MD run starting from the crystallographic HOD-QND structure^14^ revealed only limited deviations from the experimental model (Figure S6). To assess if a possible correlation exists between S101 rotamer preference and O_2_ stability at the *R*-site we then performed multiple independent simulations in which a single O_2_ molecule was positioned at this location underneath the substrate and monitored its residence times with S101 restrained either to its *trans* or *plus* rotamer. A total of 50 simulations were carried out for each rotamer. We find that the *trans* rotamer stabilizes O_2_ at the *R*-site better than the *plus* rotamer with mean residence times of 2.61±0.34 ns and 0.55±0.08 ns, respectively (Figure 3C). Also, a wide time distribution is observed when S101 is restrained to the *trans* rotamer, including times up to 8 ns, whilst times are short and tightly distributed for the *plus* rotamer. Alanine substitution (S101A) results in a behavior like S101*^t^* (mean±s.e.m. of 3.95±0.40 ns) albeit with generally longer residence times. We have also estimated residence times (*k*_off_^-1^) from the numerical fitting of the cumulative time distribution of individual events that is expected to follow Poisson statistics. This procedure gives similar *k*_off_^-1^(O_2_) values (Figure S7).

Next, we carried out 30 more independent simulations and performed a similar analysis for a H_2_O molecule located at the *R*-site (Figure 3D). As expected, water is retained at this location longer than O_2_. However, whilst for *plus*-restrained S101, water was observed leaving the *R*-site only twice within a cutoff of 50 ns (6.7% escapes, with individual residence times of 33.1 and 47.5 ns), in the case of *trans*-restrained S101 water stability is significantly reduced (96.6% escapes, mean±s.e.m. of 13±2 ns). In the case of the S101A substitution, H_2_O stability is somewhat intermediate between the behavior promoted by the two S101 rotamers (67% escapes) with a large time distribution. Numerical fitting gives a *k*_off_^-1^(H_2_O) value of approximately 15 ns for S101*^t^* (Figure S7) whilst it is greater than 50 ns for both S101*^t^* and S101A.

Overall, our analysis suggests a role for S101 as a modulator of O_2_/H_2_O stability at the *R*-site. Transient O_2_ binding within the HOD architecture favors the switch to the *trans* rotamer which, in turn, promotes both O_2_ stabilization and H_2_O destabilization at the *R*-site. The S101A replacement at the nucleophilic elbow appears to be a useful strategy to further stabilize O_2_ at the *R*-site at the cost, however, of less effective water clearing.

### Normoxic and hyperoxic HOD^H251A^-MQO structures validate S101 as an O_2_/H_2_O modulator

To visualize O_2_ at the *R*-site, we turned to pressurization experiments in the crystal state using the ‘soak-and-freeze’ method that allows crystal cryocooling under pressure.^27^ This is a significant advantage over methods that require pressure release before cryocooling as it minimizes the escape of gas molecules that often have high *k*_off_ rates.

Initially, we employed the near-inactive HOD^H251A^ variant that under normoxic conditions affords the visualization of a stable HOD^H251A^-QND complex.^13^ However, O_2_-pressurization of the complex (40 bar O_2_ for 2 minutes) leads to the complete conversion of the bound QND into the product (see Supplementary Information and Figure S8). This clearly indicates that in the crystal state under hyperoxic conditions, O_2_ can reach the *R*-site in the preformed E-S complex. To prevent turnover, we next synthesized the substrate analogue 2-methyl-quinolin-4(1*H*)-one (MQO). MQO, is the non-reactive molecule structurally closest to the natural substrate, in which the hydroxyl group at position 3 is replaced by the –H substituent. As crystals of HOD^H251A^ typically diffract better than those of wtHOD we solved the X-ray structure of this variant in complex MQO under normoxic and hyperoxic conditions at the 2.0 Å and 2.1 Å, resolution respectively (Table S1). MQO binds in the active site similarly to QND indicating that the lack of the 3OH group does not result in substantial changes. MQO is held in place by a single H-bond between its NH group and the carbonyl oxygen of W36 at 2.8 Å. Several residues contributed both by the core domain (Gly35, Trp36, Cys37, His38, His100, Ser101, His102, Gly103, Phe252) and the cap domain (Phe136, Leu140, Leu143, Leu156, Trp160, Met177, Trp185, Ser188, Gly189, Ile192) further stabilize the ligand with hydrophobic interactions (Figure 3E). A water molecule (W1) is present at full occupancy at the *R*-site underneath the A-ring of MQO at 2.8 Å from its molecular plane. It is stabilized by an interaction with the main chain amide of W36 and the side chain of S101 which is observed in double conformation in both molecules present in the a.u. with *trans* and *plus* rotamers refining at occupancies of 0.60 and 0.40, respectively. W1 is H-bonded to S101*^p^*at 2.7 Å from its OG atom, whilst S101*^t^*(OG) is more than 3.8 Å away.

We next inspected electron density maps following the O_2_-pressurization experiment (40 bar O_2_ for 2 minutes) (Figure 3F). These do not reveal changes that we could positively ascribe to O_2_ bound at the *R*-site. Instead of the elongated electron-rich density observed for bound dioxygen in other systems,^27, 28^ O_2_-pressurization led to a decrease in electron density at the *R*-site compared to normoxic conditions. Occupancy refinement of a water molecule at this location gives values of 0.67 and 0.48 in the two independent molecules present in the a.u. Remarkably, S101 shifts completely to the *trans* rotamer (χ1 values of 180.08 and 180.66 degrees) with no indication of the alternative *plus* rotamer observed under normoxic conditions.

Overall, these observations indicate that O_2_ most likely interferes with H_2_O binding at the *R*-site and although disorder negates its positive identification it nevertheless leads to the shift of S101 to the *trans* rotamer that, in agreement with the simulations, appears to be positively correlated with the presence of dioxygen at the *R*-site.

### O_2_ access to the *R*-site in the pre-formed E-S complex

As *in crystallo* O_2_ pressurization promoted turnover of the HOD^H251A^-QND complex we employed again MD simulations to gain an understanding of possible access O_2_ routes to the *R*-site in the pre-formed E-S complex. We performed a total of 80 independent 200-ns standard MD runs either in the presence of ten O_2_ molecules placed in a box of approximately 74×74×74 Å^3^ (OXY10-S-MD runs) or using a single O_2_ molecule placed initially near W37 and constrained within a sphere of 17-Å radius from S101 (OXY1-S-MD runs). The latter condition decreases the sampling space while still allowing O_2_ to diffuse outside the protein.

The simulations reveal that O_2_ entry at *R*-site is not a frequent event. Out of the 80 standard MD simulations we observed spontaneous O_2_ entry at the *R*-site, only once in either OXY10-S-MD or OXY1-S-MD runs. These events occurred when S101 was restrained to its *trans* rotamer. O_2_ access to the *R*-site followed the same trajectory in both productive runs (Figure 4A). Dioxygen entered the ABH-fold near P132 (cluster I in Figure 4A) and reached the *R*-site (cluster II) taking advantage of a hydrophobic path lined by residues belonging to α_CAP_1 (F136 and L140) and α_CAP_3 (I192 and G189). O_2_ then escaped from the *R*-site highlighting clusters III and IV near the nucleophilic elbow. These were recurrently visited prior to reaching clusters V and VI close to W36 and V71, respectively. Cluster V matches very well a high-probability O_2_ pocket seen in MD simulations in the absence of the substrate.

**Figure 4.**
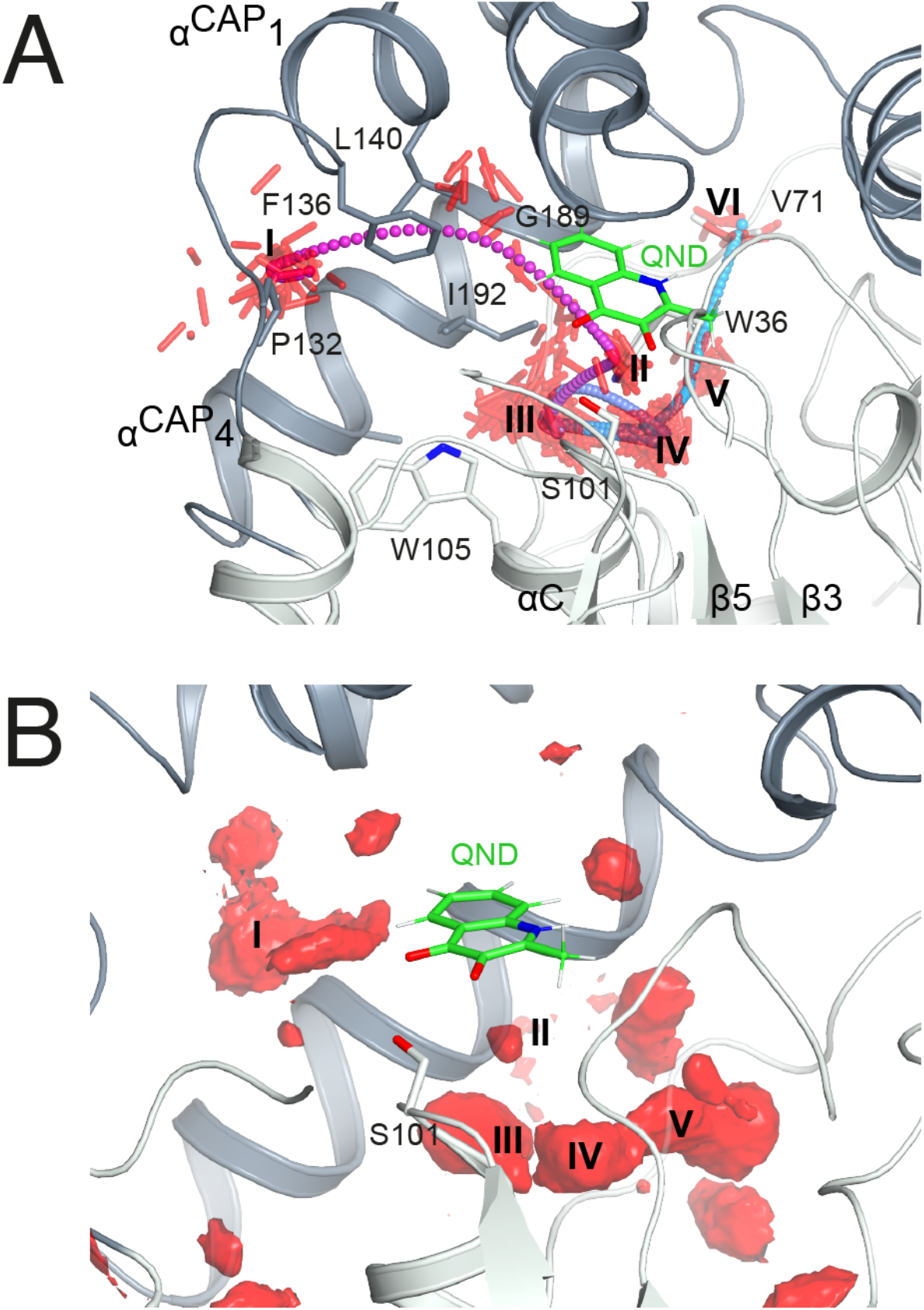
O_2_ access to the *R*-site in the E-S complex. (A) Dioxygen trajectory extracted from a standard MD simulation. Roman numbers indicate clusters of dioxygen molecules, shown as red sticks, sampled every 50 ps. The O_2_ trajectory, highlighted by small spheres colored using a cyan-magenta gradient (entry-exit), shows that dioxygen reaches the *R*-site (cluster II) from cluster I, and exits the ABH-fold via clusters (III-VI) as described in the main text. (B) regions of high O_2_ occupancy probability (isosurface representation in red) obtained reconstructing the free energy in the Cartesian space from metadynamics simulations. Roman numbers indicate the same clusters of panel A.

In a second set of calculations, we employed biased metadynamics simulations with a single O_2_ molecule placed in the bulk and free to visit any location inside the box (OXY1-S-metaD runs). The use of a bias (see Methods section) allows to accelerate the sampling of the entire 3D Cartesian space with simulations restricted to 1.8 μs each. OXY1-S-metaD runs highlight the same clusters I-V seen in the standard simulations in addition to a few additional clusters (Figure 4B). Energy calculations confirm a lower barrier for O_2_ entry when S101 is in the *trans* rotamer (4.1±1.1 kcal/mol) compared to the *plus* rotamer (6.5±1.5 kcal/mol). This corresponds roughly to a five-fold increased probability in the case of the former.

### Visualization of O_2_ at the *R*-site

As the simulations suggested that the S101A substitution decreases *k*_off_(O_2_,), we then solved the structure of HOD^S101A^ in complex with MQO analogue under normoxic and hyperoxic conditions at 2.0 Å resolution (Table S1). The HOD^S101A^ variant was also considered interesting as the AqdC ABH-fold dioxygenase that catalyzes a chemical reaction identical to HOD naturally possesses an alanine instead of a serine at the nucleophilic elbow (hydrophobic elbow).

MQO binds to HOD^S101A^ as observed with HOD^H251A^. Although the S101A replacement causes the loss of the H-bond between S101(OG) and MQO(O4) this is compensated by the interaction with H251(NE2) resulting in an essentially identical binding mode. Under normoxic conditions, electron density at the *R*-site of HOD^S101A^ is consistent with the presence of a water molecule (Figure S9). However, in contrast with what was observed for the HOD^H251A^-MQO complex, following O_2_-pressurization (40 bar O_2_ for 2 minutes) Fourier difference maps revealed a strong elongated peak at the *R*-site of one of the two HOD^S101A^ chains in the asymmetric unit (Figure 5A). This peak ranked third overall for height (+7.30) with the first two corresponding to sodium ions. Modelling of this peak as an O_2_ molecule did not result in negative difference density and occupancy refinement converged to unity with atomic displacement parameters for both oxygen atoms consistent with those of the surrounding atoms (average *B*-value MQO = 21.3 Å^2^, *B*-value O_2_(O1) = 20.5 Å^2^ *B*-value O_2_(O2) = 19.7 Å^2^). Moreover, omit maps for the individual oxygen atoms produced difference Fourier peaks, supporting the assignment as O_2_ (insets Figure 5A). The equivalent peak in the other protein chain was less prominent and more spherical in shape. We have modelled this peak also as O_2_ and occupancy refinement converged at 0.80 with an average *B*-value of 30.3 Å^2^. This value is marginally higher than that of the MQO molecule in its proximity (average *B*-value = 21.9 Å^2^) suggesting higher rotational disorder.

**Figure 5.**
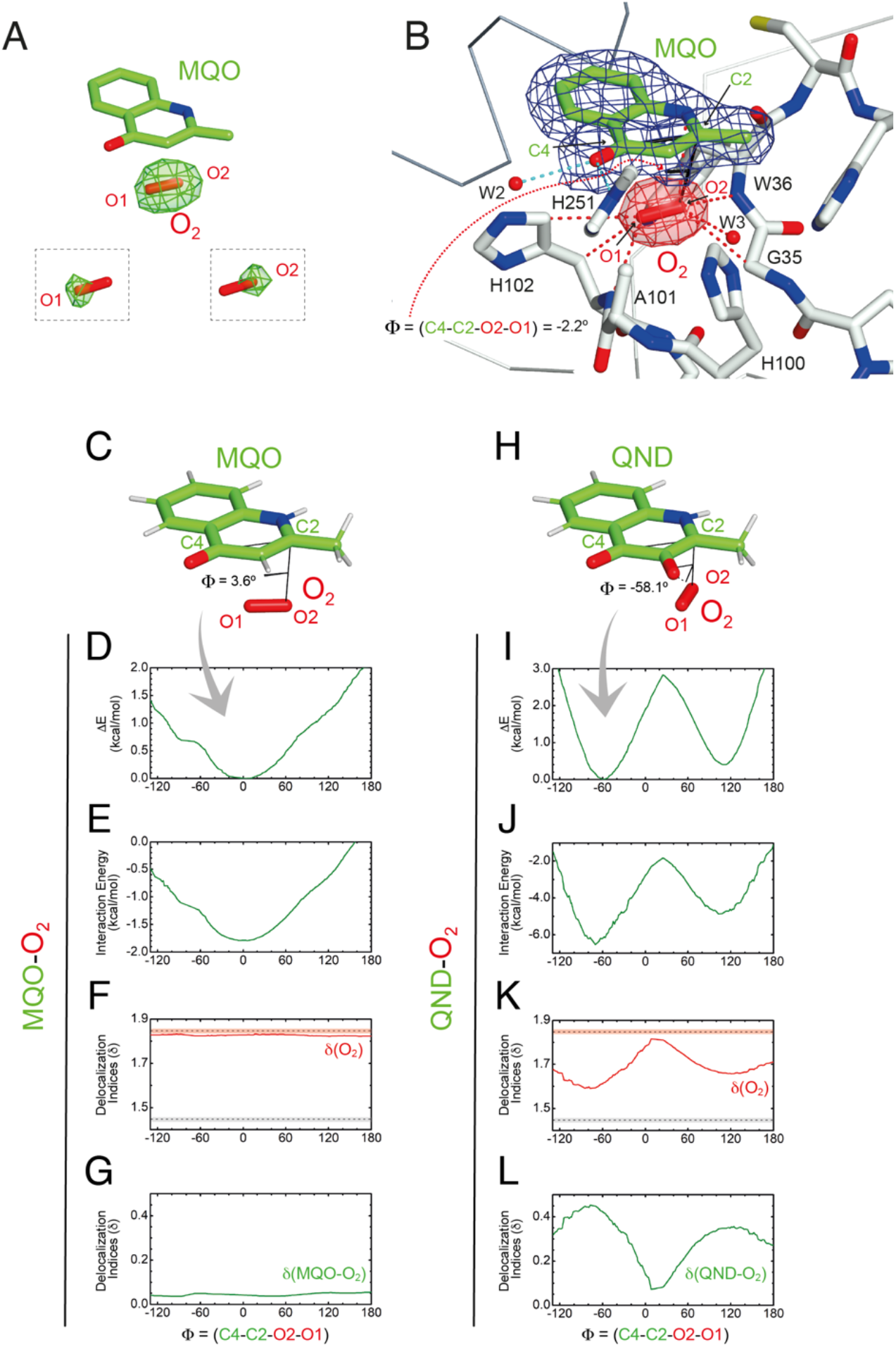
Visualization of dioxygen at the *R*-site and QM validation. (A) Following O_2_ pressurization (40 bar for 2 minutes) of the HOD^S101A^-MQO complex, *mF*_o_-*DF*_c_ Fourier difference maps at 2.0 Å-resolution (in green at the +3.0σ contour level) reveal a strong elongated peak at the *R*-site underneath the MQO ligand that is consistent with dioxygen. O_2_ from the refined model is shown for reference as a red stick. The insets show the *mF*_o_-*DF*_c_ electron density map (+3σ) with O1 and O2 atoms selectively removed from crystallographic refinement. (B) 2*mF*_o_-*DF*_c_ electron density of the HOD^S101A^-MQO-O_2_ complex. Electron density maps are shown at the +1.0σ level as chicken-wire representation for MQO (blue) and O_2_ at the *R*-site. The latter is displayed in red for clarity. The dihedral angle Φ=(C4-C2-O2-O1) in the refined crystallographic model is -2.2°. (C-L) QM geometry restrained optimization and delocalization indices (δ) for the MQO-O_2_ (C-G) and QND-O_2_ (H-L) complexes as a function of the dihedral angle Φ. (C,H) stick representations of the minimized structures, (D,I) total energy, (E,J) interaction energy-only, (F,K) δ for O_2_, (G) sum of the pair-8 between the ligand and O_2_ for the MQO-O_2_ and the QND-O_2_ complexes, respectively. In (H,J) δ values for isolated O_2_ (dashed red line) and the aromatic C-C bond in benzene (dashed grey line) are given for reference. Experiment and theory are in excellent agreement that the MQO-O_2_ complex displays an energy minimum close to Φ=0°.

Using the fully occupied O_2_ molecule as a reference, we find that dioxygen binds underneath the *A*-ring of MQO approximately parallel to its plane at an average distance of 3.2 Å (Figure 5B). The ligand’s atoms closest to O_2_(O2) and O_2_(O1) are C2 at 3.1 Å and C4 at 3.4 Å, respectively, and dioxygen further interacts with the main chain amide groups of W36 and H102 as well as with the side chain of the latter residue. Next to the *R*-site, a water molecule (W3) is also present at H-bond distance from O1 (2.7 Å) stabilized by the side chain of H100 that is flipped compared to the HOD^H251A^ variant. W3 and H100 flipping are also seen under normoxic conditions in two of the four molecules present the a.u.. The O_2_ molecular axis is roughly parallel the MQO(C2-C4) direction as quantified by the dihedral angle Φ(C4-C2-O2-O1) of -2.2°. Thus, in the HOD^S101A^-MQO-O_2_ ternary complex, dioxygen appears to preferentially adopt an orientation that mimics that of the endoperoxide during the catalytic cycle (step **4** in Figure 1B).

### The geometry and electronic properties of the ligand-O_2_ complex depend on the ligand’s charge

To seek further insight into the geometry and electronic properties of the QND-O_2_ and MQO-O_2_ complexes, we performed *ab initio* quantum mechanics (QM) calculations. Initially, we considered two systems constituted only by the ligand (either QND or MQO) and O_2_. As QND is deprotonated by the His-Asp dyad, we assumed that the QND-O_2_ system bears a single negative charge whilst MQO-O_2_ is neutral. In these calculations, we restrained the sum of the (C2-O1) and (C4-O2) distances to the crystallographic values. However, around the equilibrium geometry, our results are insensitive to small changes in the restraint value (Figure S10).

The total energy of the complexes as a function of Φ is shown in Figure 5D,I. Despite the structural similarity between QND and MQO, the QM calculations revealed that the orientation of O_2_ with respect to the ligand is different. Whilst in the MQO-O_2_ complex, dioxygen orients its axis along the MQO(C2-C4) direction with a dihedral angle Φ of 3.6° (Figure 5C) in excellent agreement with our crystallographic structure, in the QND-O_2_ complex, O_2_ is aligned with the QND(N1-C3) bond at an angle of about -58.1° (Figure 5H). We observe that an essentially identical angular dependence is observed for the “interaction energy”-only component of the total energy (Figure 5E,J) indicating that molecular distortions are not important in defining the equilibrium geometry of these systems. Energy decomposition analysis allowed to further investigate the contribution of the different terms (Pauli repulsion, charge transfer, electrostatic, dispersion, and polarization) to the interaction energy (Figure S11).^29, 30^ For the MQO-O_2_ complex, the most stable orientation is that which minimizes the Pauli repulsive energy between the electronic clouds of MQO and O_2_, that is, in this case, associated with π-π stacking. Differently, for the QND-O_2_ complex, the equilibrium geometry is that which maximizes the charge-transfer from the QND anion to O_2_. We hypothesized that the difference in the relative O_2_ orientation between the two complexes is caused by the negative charge of QND. This was confirmed by calculations for the protonated (neutral) QNDH-O_2_ complex, that mirror closely those obtained for MQO-O_2_ (Figure S12).

Next, we analysed the electron density shared between the ligand and O_2_ using delocalization indices (δ), which are intimately related to bond order.^31^ We calculated these values as a function of Φ for dioxygen δ(O1-O2), and for the sum of all the pair-δ values between the ligand and O_2_, δ(QND-O_2_) and δ(MQO-O_2_). For the MQO-O_2_ complex, we find that independently of Φ, δ(MQO-O_2_) is close to zero whilst δ(O1-O2) is close to the value for isolated O_2,_ δ(O1-O2)=1.84 (Figure 5F,K). This indicates that there is essentially no electron density transfer between MQO and O_2_. For the QND-O_2_ complex, both δ(QND-O_2_) and δ(O1-O2) are no longer independent of Φ. Moreover, at the equilibrium geometry, where charge-transfer is maximized, δ(QND-O_2_) has a maximum at ∼0.35 while δ(O1-O2) has a minimum at ∼1.67, which is intermediate between the value for isolated O_2_ and that of an aromatic C-C bond (Figure 5G,L). In this complex therefore electron density is shared between QND and O_2_ that results from the electron transfer from the π cloud of QND to the π* orbital of O_2_, decreasing its bond strength.

To test the effect of the protein matrix we have repeated the same analysis above using a QM/MM model in which the QM region includes the substrate, O_2_ and key active site residues (Figure S13A,E). In these calculations that explicitly include the active site environment, no restraints were employed. These more sophisticated calculations do not alter our previous conclusions and the results are entirely consistent with that obtained for the isolated complexes (Figure S13B,F). We note, however, that for MQO-O_2_, the energy difference associated with O_2_ rotation is smaller, with a barrier height lower than 0.5 kcal/mol, which could be easily overcome at thermal energies. This flat energy profile allows therefore rotational averaging which likely contributes to an increase in the sphericity of O_2_ electron density at the *R*-site as observed in one of the two active sites. For the QND-O_2_ complex, the barrier is about 2 kcal/mol, like that obtained from the isolated complex. Analysis of the delocalization indices also produced results like those obtained for the isolated complexes (Figure S13C,D,G,H).

### Reaction mechanism

To investigate the reaction mechanism, we calculated the QM/MM energy profile along the reaction coordinate α (Figure 6A). The reaction can be conveniently rationalized as composed of two stages: *i*) an intersystem crossing (ISC) stage in which the system ‘hops’ from the triplet to the singlet state with the formation of the C2-peroxide (**2**) and *ii*) the CO-release stage, in which the peroxide breaks down via the endoperoxide (**3** and **4**) leading to products (**5**). For the ISC stage, the energy profile was calculated as a function of the energy difference between the singlet and triplet states (Figure S14) and then represented as a function of the reaction coordinate α as shown in Figure 6A.

**Figure 6.**
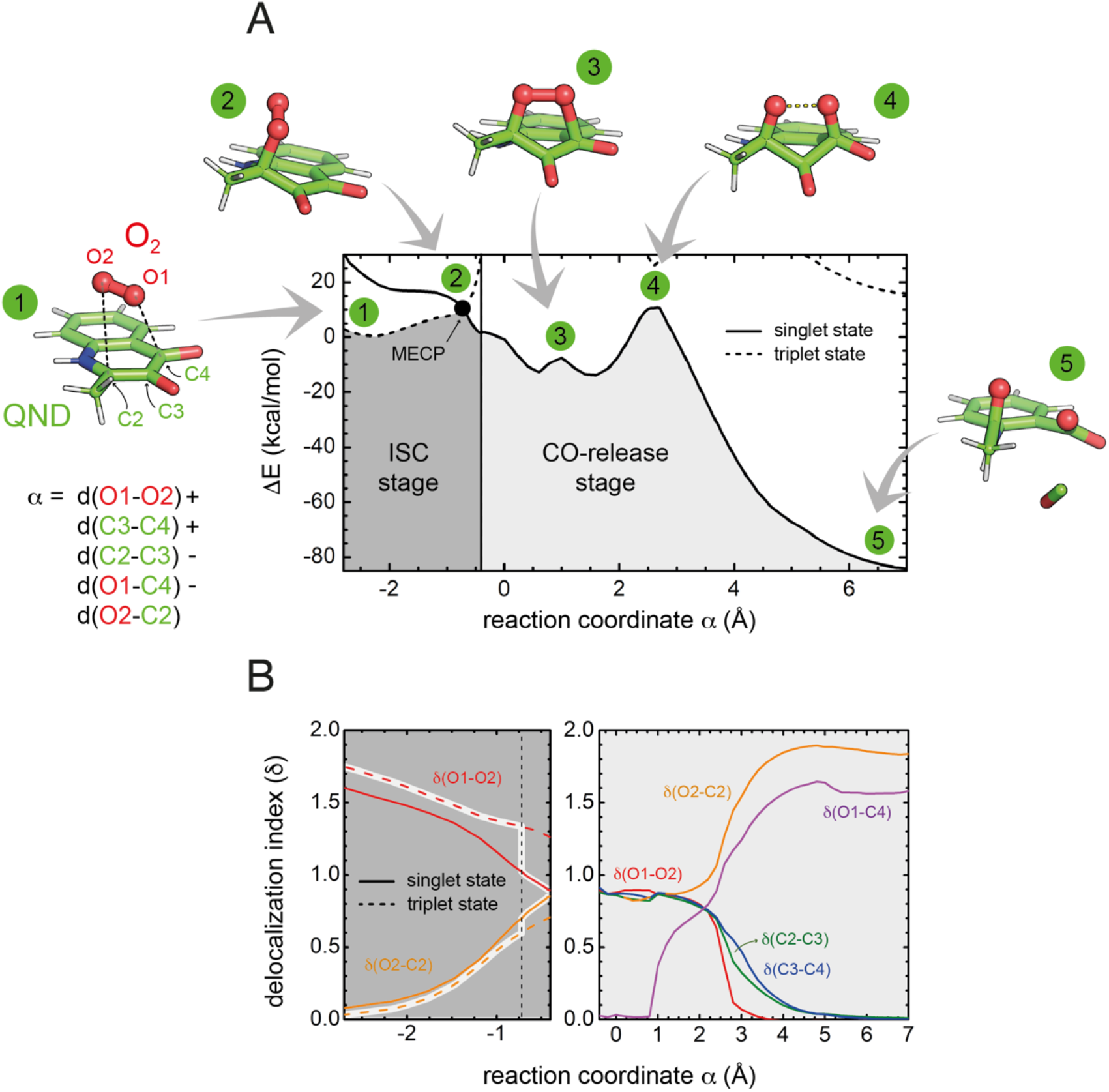
Energetics and charge transfer along the reaction path. (A) Energy profile calculated at the M06-2X/6-31+G(d,p) level of theory as a function of the reaction coordinate α. The lowest energy singlet and triplet states are shown as a continuous and dashed lines, respectively. Numbered key stages of the reaction are visualized as ball-and-stick representations. In all of them, the protein is not drawn for simplicity. The vertical line (at α = -0.4) distinguishes the two reaction stages. The left-hand region defines the intersystem crossing (ISC) stage, in which the system switches from the triplet to the singlet state leading to the formation of the peroxide through the minimum energy crossing point (MECP, highlighted by a black dot). The CO-release stage on the singlet potential energy surface is strongly exothermic. (B) Evolution of the delocalization indices (δ). The left panel shows δ indices for the ISC stage where only δ(O1-O2) and δ(O2-C2) change significantly. The DIs are shown for the triplet (dashed lines) and singlet (solid line) states, and those corresponding to the most stable state are highlighted. The vertical dashed line represents the position of the MECP. Differences between δ values for the singlet and triplet states at MECP indicate that ISC is associated with a significant transfer of electronic density from the substrate to O_2_. In the right-hand side panel, δ indices are shown only for the singlet state, as the triplet state is very high in energy.

Reactants (**1**) are stable in the triplet state arising from the combination of the electronic ground state of O_2_(^3^Σ_g_^-^), and the ground state of QND (singlet, all electrons are paired). However, as QND and O_2_ move closer to each other, the energy of the system increases and any attempt to optimize the C2-peroxide in the triplet state led back to the reactants. On the contrary, formation of the peroxide in the singlet state proceeds barrierless leading to a stable moiety. Analysis of the singly occupied molecular orbitals (Figure S15) suggests that stabilization and destabilization of the singlet state and of the triplet state, respectively, originate from the combination of one π* orbital of O_2_ with the π cloud of QND. This leads to an orbital with antibonding character along the C2-O2 and O1-O2 bonds, which is singly occupied in the triplet state and unoccupied in the singlet state. Singlet and triplet states must cross at some point along the reaction path and the minimum energy crossing point (MECP, highlighted by a black dot in Figure 6A), acts as the effective transition state. Our calculations locate the MECP (**2**) at 10.2 kcal/mol above the energy of the activated QND-O_2_ complex (**1**). At the MECP, d(C2-O2) is 1.62 Å, and QND is no longer planar. To get further insight on the ISC stage, we evaluated how δ(O2-C2) and δ(O1-O2) change along the reaction path in both the singlet and triplet states (left-hand panel in Figure 6B). We observed that shortening of d(C2-O2), results in an increase of the electronic density shared between the substrate and dioxygen, which leads to the weakening of the O_2_ bond, a consequence of the larger charge transfer from QND to O_2_. Moreover, at every point along the reaction path, charge transfer is stronger in the singlet than in the triplet state, so the spin-flip involves a significant transfer of electronic density between QND and O_2_.

Once the peroxide is formed, the next stage of the reaction proceeds via the formation of the endoperoxide (**3**) and subsequent rupture of the C2-C3 and C3-C4 bonds followed by CO release that is very exothermic. The process is again well described by the delocalization indices (right-hand panel in Figure 6B). Once the system is in the singlet state, the peroxide attacks the carbonyl group establishing the C4-O1 bond of endoperoxide. At the barrier (**4**), concerted breaking of the endoperoxide and CO release takes place. As shown in Figure 6B, the energy of the transition state for the CO-release stage lies 10.8 kcal/mol above the reactants, which is slightly above the energy of the MECP. However, to compare the barrier heights it is necessary to correct the ISC energy barrier for the probability of hopping from the triplet to the singlet state,^32^ which depends on the value of the spin-orbit coupling (SOC). At the MECP, we calculated a SOC of 22 cm^-1^, leading to a swapping probability of 0.005, i.e. an increase of 3.1 kcal/mol of the barrier (see supplementary material for further details). Therefore, the ISC stage is the rate limiting step of the overall process.

We also considered the possibility of a direct electron transfer within the complex by calculating the energy difference between [^3^O_2_ + ^1^QND^-^] and [^2^O_2_^●^ + ^2^QND^●^] at the reactants’ asymptote, based on separate calculations for the donor and the acceptor. We obtained a value of 17 kcal/mol, which is significantly larger than the energy of the MECP (even when corrected by the swapping probability). This result is compatible with our findings that, along the reaction path, the energy of the singlet state monotonically decreases. Although such an endothermicity does not allow to completely rule-out the possibility of a direct electron transfer,^17^ our calculations predict that a scenario where spin-flipping is more favorable. Consistent with this, no significant amounts of radical species were detected in spin-trapping experiments with wild-type HOD.^33^

## Concluding remarks

The “great oxidation event” of 2.4 billion years ago led to the permanent accumulation of O_2_ in the atmosphere thus exerting evolutionary pressure on enzyme systems that could take advantage of such a strong oxidant. A very recent KEGG (Kyoto Encyclopedia of Genes and Genomes) analysis of the 136 Pfam families of all known O_2_-dependent enzymes, found that only ∼60% of these can be classified as having a function primarily related to O_2_ whilst the remaining ∼40% represent families featuring only sporadic O_2_ utilizers.^1^ The latter group has therefore likely evolved O_2_-metabolizing capabilities from a different set of original catalytic competences. Hydrolases, particularly metallo-β-lactamases and those of the ABH-fold, appear to be the most common progenitors.^1^ ABH-fold dioxygenases are particularly intriguing as they accomplish O_2_ redox chemistry without external cofactors.

Here, using HOD as a paradigm for the growing family ABH-fold dioxygenases, we have shown how evolution has taken advantage of multiple structural elements of the ABH-fold to enable the switch from hydrolytic to oxygenolytic reactions. A key requirement for such catalytic repositioning is the ability to host O_2_ sufficiently close to its *N*-heteroaromatic substrate for the reaction to take place. Using *in crystallo* O_2_-pressure freeze-trapping we have visualized the ternary complex with a substrate analog at high resolution thus providing conclusive evidence that the oxyanion hole fulfills the role of O_2_ receptor. Computational analyses of dioxygen diffusion pathways in various non-ABH-fold enzyme systems including flavoenzymes,^34^ the nonheme iron-containing 12/15-lipoxygenase,^35^, and also the metal-independent DpgC dioxygenase ^19^ have uncovered some common themes in O_2_ access strategies, often involving a network of transient pathways that are mostly lined with hydrophobic residues. In HOD, we have identified a set of pockets that provide a route for O_2_ access to the oxyanion hole in the E-S complex. However, compared to other systems O_2_ entry appears a much rarer event arguing in favor of additional strategies to increase the local O_2_ concentration. In urate oxidase, a hydrophobic cavity adjacent to the active site has been suggested to act as transient O_2_ reservoir.^36^ We propose that in HOD, and likely in all ABH-fold dioxygenases that operate on similar organic substrates, the active site itself acts as a temporary O_2_ binding site. Transient O_2_ storage at the xenon-validated *B*-site that we have identified overlapping with the hydrophobic portion of the *N*-heteroaromatic binding site represents a convenient solution to the problem O_2_ localization as one might envisage a mechanism of displacement that upon substrate binding transfers O_2_ from the *B*-site to the *R*-site that is only about 5 Å away thus organizing both reactants for catalysis.

ABH-fold enzymes often employ their typical Ser-His-Asp triad to catalyze reactions following an esterase-type mechanism. Departures from this have however been observed in some non-hydrolytic reactions. *Manihot esculenta* (cassava) and *Hevea brasiliensis* (rubber tree) hydroxynitrile lyases that break the C–C bond in cyanohydrin although featuring the classical Ser-His-Asp triad, they use their active site serine as a general base. Even more dramatic is tomato methyl ketone (MK) synthase (MKS1) involved in MK 2-tridecanone synthesis that lacks the canonical triad whilst retaining a conserved histidine residue that acts as the catalytic base with a neighboring threonine providing the H-bond bond pattern required for the decarboxylation step of β-keto acid substrates.^37^ For oxygenation, the ABH-fold catalytic machinery is also employed in an unconventional manner and substrates’ activation by deprotonation of their 3-hydroxyl group only requires the His-Asp subset (H251-D126 in HOD) that acts as a general base.^13, 14^ Although HOD’s serine residue (S101) has been shown to retain nucleophilic properties in some hydrolase-like reactions,^38^ this capability is irrelevant for oxygenation.^14^ Instead, our results indicate that in addition to a contribution in the stabilization of the organic reactant,^14^ S101 plays a role as O_2_*/*H_2_O modulator with its *trans* rotamer promoting both O_2_ stabilization and H_2_O destabilization in the oxyanion hole. Substitution of the cryptic serine nucleophile with an alanine, a replacement naturally present in some ABH-fold dioxygenases that further strengthens the argument of nucleophile dispensability, also promotes O_2_ stabilization at the catalytic site. In HOD, S101A replacement has however a negative impact on the *K*_M_ value for its organic substrate (60-fold increase compared to wild-type HOD) whilst *k*_cat_ is not affected. ^14^ This suggests that the evolutionary choice of the specific amino acid at the ‘nucleophile elbow’ is likely the result of a combination of factors aimed at maximizing the stability of organic substrate and O_2_ whilst achieving destabilization of H_2_O.

Deprotonation of the substrate is a common step in cofactor-independent oxygenation^11, 39^ and our QM and QM/MM analyses reveal that the charge of the substrate determines the geometry of the O_2_-substrate complex. Cofactorless addition of O_2_ to QND requires the swapping from the triplet to the singlet state, a “spin-forbidden” process. The degree of spin-forbiddenness is determined by the magnitude of the SOC term which correlates with the energy difference between the two π-orbitals of O_2_. When QND is deprotonated, the geometry of the O_2_-QND complex is that which maximizes electron density transfer from QND (π) to O2 (π*) orbitals, thus perturbing the π-symmetry of O_2_ and leading to values of SOC large enough to make the reaction feasible,^40^ even in the absence of any metal cofactor. For spin-forbidden reactions, the MECP acts as the main dynamical bottleneck for the reaction,^32^ and we calculated that the additional burden caused by the spin-forbidden character of the reaction is 3.1 kcal/mol. Similarly to what was obtained for DpgC^41^, vitamin K-dependent glutamate carboxylase,^42^ or nogalamycin monoxygenase,^43^ O_2_ is not protonated at the MECP, in contrast to what was obtained for p-hydroxyphenylacetate hydroxylase, a flavin-dependent monooxygenase.^44^

To summarize, evolution has repurposed the ABH-fold architecture and its simple catalytic machinery to accomplish metal-independent oxygenation. This is achieved by dioxygen entrapment at the oxyanion hole in proximity of the organic ligand with its stabilization and concomitant H_2_O destabilization modulated by the nucleophile/hydrophobic elbow residue (Ser/Ala). Substrate deprotonation mediated by the His-Asp subset of the catalytic triad elicits a ligand-O_2_ geometry that maximizes electron transfer such that the spin-forbiddenness of the direct reaction is relaxed.

## Acknowledgments

SB was supported by the United Kingdom Biotechnology and Biological Sciences Research Council (BBSRC) grant BB/I020411/1 awarded to RAS. PGJ gratefully acknowledges grant PID2020-113147GA-I00 funded by the Spanish Ministry of Science and Innovation MCIN/AEI/10.13039/501100011033. PGJ and SGG acknowledge funding from the Fundación Salamanca City of Culture and Knowledge (programme for attracting scientific talents to Salamanca). RAS acknowledges the Regione Autonoma della Sardegna (decision n. 56/21) for sponsoring his visiting professorship at the University of Cagliari. We thank Professor Thierry Prangé (Université Paris Cité) for his help with the crystallographic room temperature pressurization setup.

## Author contributions

RAS conceived the project. SB performed the experimental work. SGG, MC, and PGJ carried out the computational work. PvdL and PC contributed to the crystal pressurization experiments. SB, SGG, MC, PGJ, and RAS analyzed data. The manuscript was written by RAS with contributions from MC and PGJ and commented on by all authors.

## Supplementary Information for

### DNA techniques, protein expression, and protein purification

For protein overexpression in *E. coli*, the target gene was inserted into the pQE30 vector (Qiagen) at the BamHI and SalI restriction sites. An uncleavable N-terminal His_6_ fusion tag of sequence MRGSHHHHHHGS was added to the gene product to facilitate protein purification. As wild-type HOD tends to dimerize due to oxidative formation of an intermolecular disulfide bridge, we took advantage of the C69S substitution that abrogates this problem.^1^ The HOD C69S variant (HOD^C69S^) has catalytic properties that are identical to those of wild-type HOD. All work described here was carried out in this background. For simplicity, throughout the text, we refer to His_6_-HOD^C69S^ simply as HOD. Site-directed mutagenesis to generate all HOD variants was performed using standard techniques followed by gene sequencing for validation. Protein over-expression was accomplished in *E. coli* M15[pREP4] cells at 28 °C. Purification of all HOD variants was performed using a combination of chromatographic techniques as described previously. The substrate 1-*H*-3-hydroxy-4-oxoquinaldine (QND) was a kind gift of Professor S. Fetzner (University of Münster, Germany). The substrate analogue 2-methyl-quinolin-4(1*H*)-one (MQO) was commercially synthesized by AF ChemPharm, Sheffield, UK.

### X-ray crystallography

Crystals of HOD, HOD^H251A^, HOD^S101A^ exhibiting variable degree of twinning and belonging to space group *P*2_1_2_1_2_1_ or *P*4_1_ were obtained by the vapor-diffusion method at 18 °C by mixing equal volumes of protein solution (150 mg/ml in 20 mM Tris-HCl,100 mM NaCl, 2 mM EDTA, 1mM DTT, pH 7.5) and a reservoir solution containing 1.65 M Na/K tartrate and 100 mM Hepes, pH 7.0. Complexes with the organic compounds were obtained by soaking crystals in a reservoir solution enriched by a 2 mM solution of the appropriate ligand for typically around six hours. Noble gas derivatization was achieved by pressurizing at 30 bar a capillary-mounted HOD crystal in pure Xe atmosphere. Pressure was maintained for the entire duration of the room-temperature diffraction experiment. For dioxygen pressurization experiments, we employed the methodology dubbed ‘soak-and-freeze’.^2^ Crystals were mounted in specific sample high pressure-supports (SPINE standard pins with a capillary) and exposed to 40 bar O_2_ for two minutes at ambient temperature prior to cryo-cooling in liquid N_2_ whilst still under pressure. Except for the Xe pressurization measurement, all data collections were performed at 100K with crystals cryo-protected using the reservoir condition spiked with 20% glycerol (v/v). Complete data sets were measured at beam lines I02, I04-1, I24 of Diamond Light Source (Didcot, United Kingdom) or ID29, BM30A of the European Synchrotron Radiation Facility (Grenoble, France). Data processing was carried out using the *xia2* pipeline^3^ and crystal structures solved either by the molecular replacement technique using *Phaser^4^* or by difference Fourier methods. Model refinement was achieved with the *Refmac*5^5^ package. Model building was performed with COOT.^6^ A summary of data collection and refinement statistics are shown in Table S1.

### In crystallo turnover

In solution, the H251A replacement renders the enzyme almost inactive (*k*_cat_ of 0.0006 s^-1^ at pH 6.5 compared to 16.3 s^-1^ for wtHOD) as it lacks the functional H251-D126 dyad required for the fast substrate activation step in which its 3OH group is deprotonated.^7, 8^ Previously, we have obtained the X-ray structure of the HOD^H251A^-QND complex under normoxic conditions.^7^ The structure shows that under these conditions the substrate binds in the active site without any appreciable turnover. Despite the very limited activity of HOD^H251A^, O_2_ pressurization of HOD^H251A^-QND crystals (40 bar for 2 minutes) results in complete turnover and electron density maps at 2.0 Å show the bound N-acetylanthranilate product (Figure S8A). This experiment clearly demonstrates that O_2_ can reach the *R*-site in the preformed E-S complex promoting QND conversion into the NAA product during pressurization. Reactivity must therefore be attributed to a small quantity of the QND anion present at the equilibrium. *In crystallo* UV/Vis spectra are consistent product formation without substantial accumulation of the substrate anion (Figure S8B). The structure suggests that proton abstraction is mediated by solvent molecules that connect the 3OH group of the substrate with D126 (Figure S8A).

### Standard molecular dynamics (MD) simulations

Initial coordinates for substrate-free simulations were taken from the relevant crystallographic structure (PDB code 2WJ3). The enzyme was solvated by in a cubic box of 85×85×85 Å^3^, with water molecules placed initially 5 Å away from the protein surface whilst retaining all crystallographic water molecules. The system was initially relaxed using the NAMD package^9^ while production runs were performed with ACEMD.^10^ For the first 2 ns, restraints were applied to both Cα and Cβ protein carbon atoms and to crystallographic waters. At T = 300 K in the NPT ensemble, all restraints were gradually removed over 4 ns. After 2 ns in the NPT ensemble at 300 K and P = 1 atm, we calculated the average size of the box and we used this value for simulating the system in the NVT ensemble (L=73.67 Å, 39081 atoms). Long-range electrostatic interactions were treated with the Soft-Particle-Mesh-Ewald algorithm^11^ (64 grid points with direct cutoff at 9.0 Å, as suggested by the ACEMD software; the same cutoff holds for van der Waals interactions). We employed the Amber99SB ILDN forcefield^12^ and TIP3P^13^ for the protein and water molecules, respectively. Two independent 1 μs-long replica simulations were carried out without or with explicit O_2_ (WAT-MD and OXY-MD, respectively). For OXY-MD, ten O_2_ molecules were added to the system in the cubic box with L = 73.67 Å. O_2_ was parametrized with the LJ Amber parametrization, by adding a small charge to have a dipole of 0.121 eÅ. Normalized dioxygen distribution was calculated by sampling O_2_ positions every 50 ps onto a matrix of 0.6 × 0.6 × 0.6 Å^3^ elementary cubes and dividing the total atom count in each volume by the total number of frames.

Initial coordinates for substrate-bound simulations were taken from the anaerobic HOD-QND complex (PDB code 2WJ4) and the system was setup as described above. QND was initially considered in its protonated form at O3 (QNDH) with neutral H251 protonated on ND1. The force field for the substrate was obtained as described previously.^14^ The protonation of all other histidine residues was chosen to preserve the H-bond pattern observed in the X-ray structure. At 300 K in the NPT ensemble, we removed protein restraints over 4 ns and transferred the proton from the substrate’s O3 atom to H251(NE2) as this represents the mechanistically-relevant complex.^7^ Weak restraints on the substrate positioning were maintained throughout to prevent its escape from the active site during the runs. To study O_2_ entry in the HOD-QND complex we performed a total of 80 independent 200-ns MD runs by adding to the system either ten O_2_ molecules in the L = 73.67 Å box (OXY10-S-MD runs) or one O_2_ molecule confined in a sphere with a 17-Å radius of centered on S101 (OXY1-S-MD runs). Of the 40 OXY10-S-MD runs, 25 were carried out with a harmonic restrains on S101 to maintain its *trans* rotamer and 15 had no restraints on S101. Of the 40 OXY1-S-MD runs, 20 were carried out with a harmonic restrains on S101 to maintain its *trans* rotamer and 20 had no restraints on S101.

### Metadynamics Simulations

To investigate the oxygen diffusion within HOD, we also performed metadynamics simulations employing a bias on the O_2_ position.^15^ As coordinates to be biased we selected the position of the dioxygen center of mass in the 3D Cartesian space. As consequence of the use of absolute coordinates, we kept the protein frame in place by restraining the position of a set of Cα atoms (residues 31, 101, 112, 243) to the reference structure obtained after relaxation. We performed well-tempered metadynamics,^16^ DT= 3000 K, with bias added as Gaussians with a frequency of 1 ps. As the *R*-site cavity is rather small, we decided to work with narrow Gaussians (width=0.3 Å). For this reason, we started with a high value for the initial height (1 kcal/mol) and this value is smoothly decreased by the well-tempered algorithm during long runs. Dioxygen was confined to a sphere of 17-Å radius centered on S101. We performed three simulations of 1.8 μs each. In two replica runs (identical conditions except for initial velocities chosen randomly) S101 was restrained to its *trans* rotamer. No restraints were employed in the third one. We calculated the error on metadynamics by reweighting the positions sampled during the runs with the corresponding bias term and comparing the free energy surface with the one obtained integrating the bias potential.

### Analysis of O_2_ and H_2_O escape times from the *R*-site

Preliminary tests indicated that O_2_ residence times at the *R*-site are in the order of ns, corresponding to an energy barrier of few kcal/mol, achievable with thermal motions (*k*T = 0.6 kcal/mol at 300 K). Thus, the use of enhanced sampling techniques such as metadynamics^17^ or TPS-derived methods^18^ was not required for this analysis. As our starting point we took advantage of a protein state found in our OXY1-S-MD run in which O_2_ was found at the *R*-site. This was subjected to 200 steps of conjugate gradient minimization and then thermalised for 1 ns at 300 K before assigning new random velocities to 50 independent standard MD simulations. For each trajectory we evaluated the time spent by O_2_ at the *R*-site before it abandons this location. The escape process can be seen as a diffusion process over a barrier dividing two basins. Although there are uncertainties in the estimation of the precise time when a molecule goes over the unknown transition state for calculating the residence time,^18^ we can associate to the transition over a barrier the abrupt change in a collective variable/parameter. By plotting the distance of oxygen from the substrate geometric center (calculated using heavy atoms only), we recognized that around 6 Å there is an abrupt change of position. Thus, we defined the residence time as the time to reach in the sampled trajectories that distance, without considering the possibility to recross back the transition state, or considering the transmission coefficient equal to one. Harmonic restraints were employed to preserve S101 rotamers and the substrate position (on average) during the simulations.

Escape times (*k*_off_^-1^) are expected to follow the exponential distribution:

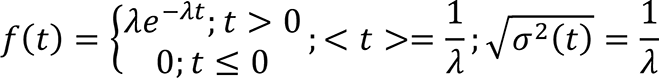

belonging to the family of Poisson statistics. From this distribution we can derive the probability that an event occurs before time *t*:

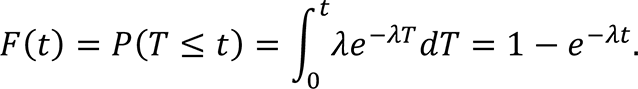

Numerically we can fit F(*t*) with the cumulative histogram of escape times to obtain the single parameter lambda that characterizes this distribution. We can also determine the average escape time using block averages. Fitting the cumulative distribution is however more appropriate because it is difficult to sample long escape times. The cumulative distributions were obtained by accumulating data with a bin width of 1 ns.

### QM calculations of the isolated ligand-O_2_ complexes

DFT calculations to determine the equilibrium geometries of the ligand-O_2_ complexes were carried out at the M062X/6-31+G(d,p) level of theory using QChem 5.2^19^ using the D3(0) dispersion correction.^20^ Since even for simple systems, such as toluene-O_2_ there are many conformers possible,^21^ we have reduced the dimensionality of the problem by restraining the sum of the (C2-O1) and (C4-O2) ligand-O_2_ distances to their crystallographic values. We find however that around the equilibrium geometry, results are quite insensitive to small changes in the restrain value (Fig. S9). This restraint was not applied in the QM/MM calculations, as the protein scaffold prevents O_2_ motion away from its pocket. Energy decomposition analysis (EDA)^22^ as implemented in QChem 5.2 was employed to calculate the interaction energy between O_2_ and QND/MQO, and its contributing terms. Delocalization indices^23^ (δ) using the partitioning Mulliken scheme^24^ were calculated using the NDELOC code,^25^ interfaced with Gaussian16.^26^. They are a measure of ‘electron sharing’ between sets of atomic centers, and their 2-order magnitudes allowed us to assign “bond orders”. To avoid a wrong partition within the Hilbert space, diffuse functions were not included in δ values calculations.

### QM/MM energy minimizations

The HOD-MQO/QND-O_2_ ternary complexes used as starting point for the QM/MM calculations were prepared from the relevant crystallographic structures using the CHARMM software^27^ and the CHARMM36 forcefield.^28^ MD simulations were performed with NAMD 2.13.^9^ The coordinates of the hydrogen atoms were generated using the standard protonation states for the titrable residues. H251 was protonated in the HOD-QND system, as this residue abstracts the 3OH proton from QND. The protein complexes were placed in the center of a cubic TIP3 water box, large enough to include the protein and at least 10 Å of solvent on all sides. KCl at a concentration of 0.15 M was added to neutralize the system, and the Particle Mesh Ewald method^29^ was used to treat the electrostatics of the periodic boundary conditions. The system was subjected to a classical MD equilibration of 10 ns at constant temperature (303.15 K) and pressure (1 atm), where to avoid O_2_ diffusion outside the active site, the position of QND/MQO and O_2_ were restrained throughout the equilibration. After equilibration, the system was trimmed to a sphere of 26 Å centered around O_2_, where the atoms further away than 18 Å were kept frozen during all QM/MM calculations. Full electrostatic embedding was adopted, and hydrogen link atoms were used to treat the QM/MM boundaries, when necessary. QM calculations were carried out at the M062X/6-31+G(d,p) level of theory using Grimme’s dispersion correction as in the gas-phase calculations. In a first set of calculations, the reactants were optimized and, subsequently, the (C4-C2-O2-O1) dihedral angle (Φ) was scanned through geometry restrained minimizations. For this study, the QM region included the substrate, O_2_, the sidechain of H251 (protonated for QND-O_2_ simulation), H102, G103, the backbone of G35, W36, C37, and three water molecules close to either O_2_ or the histidine sidechain. No restraints were applied the (C2-O1) and (C4-O2) distances. In a second set of calculations, we calculated the energy profile along the reaction path. For the CO-release stage, we used as a reaction coordinate α = d(O1-O2) + d(C3-C4) + d(C2-C3) – d(O1-C4) – d(O2-C2), which includes all bond distances that are formed and broken during the reaction. Calculations with and without the additional restraint of d(C3-C4) = d(C2-C3) were run, showing that including this restraint aids to reach the product state, without modifying the barrier heights. To ensure convergence, geometry restrained minimizations were carried out forwards and backwards. Calculations were carried out using two different QM regions, one including just the substrate (QND or MQO) and O_2_, and another also including the sidechain of H100, S101, and H251 as well as a water molecule nearby. The results were found to be relative insensitive to the size of the QM region. To describe the intersystem crossing (ISC) stage, the energy difference between the singlet and triplet states was chosen as a reaction coordinate,^30^ so the energy of the Minimum Energy Crossing Point (MECP) is that obtained obtained when the energy difference is zero. The results are shown in Fig. S13. For this system, the energy difference between the triplet and singlet states was strongly coupled with the approaching of O_2_ to QND, so it was also possible to represent the reaction pathway as a function of the composite reaction coordinate α. This is done in Figure 6 of the main text, where all the geometries for the intersystem crossing stage appear for α < -0.4 Å. For the ISC stage we only considered the small QM region, and for the sake of simplicity only results with the small QM region are shown in Figure 6.

### Calculation of the energy barrier for the intersystem crossing (ISC) step

To compare the effective energy barriers between for two reactions, one spin-allowed (adiabatic) and another spin-forbidden, the barrier height of the latter should be corrected to account for the limited swapping probability between the electronic states involved. According to the Non-Adiabatic Transition State Theory, the rate coefficient of a spin-forbidden transition should be calculated as:^31^

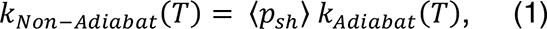

where *k_Non-Adiabat_* is the actual rate coefficient of the process, *k_Adiabat_* would be the rate coefficient of a hypothetical analogous spin-allowed reaction (just one electronic state), and 〈p_sh_〉 is the probability of spin-flipping, in this case the probability of hopping from the triplet to the singlet state at the MECP. (p_sh_) can be calculated from the Landau-Zener theory.^32, 33^ Using the double passage version of the spin-diabatic Landau-Zener formula for a given energy is,

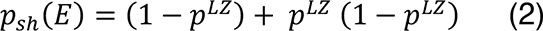

where *p*^*LZ*^ is given by:^34^

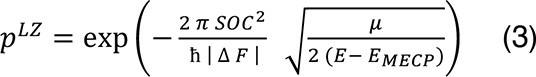

where *E* is the energy, *E_MECP_* is the energy of the MECP, μ is the reduced mass of the normal mode orthogonal to the crossing seam, | ΔF | is the norm of the difference between the gradients of the two Potential Energy Surfaces at the crossing point, and SOC is the effective spin-orbit coupling calculated at the MECP. For *E* < *E_MECP_*, *p^LZ^* = 0.

SOC was calculated at the CASPT2(22,38)/cc-pvdz level of theory using Open-MOLCAS,^35^ with the active space formed by all the valence orbitals of O1, and O2 and those related to the C2-O2 and O1-O2 bonds. The value obtained (SOC= 22 cm^-1^) was found to be barely sensitive to small changes in the active space. | ΔF | and μ where calculated following Ref. ^36^. We obtained (p_sh_) = 0.005 which leads to an increase of 3.1 kcal/mol to the barrier height.

According to our calculations the energy of the MECP was 10.2 kcal/mol above the energy of the triplet QND^-^ O_2_ complex. At the MECP, r_C2-O2_ =1.62 Å, and the QND moiety is no longer planar. The geometry at the MECP is similar to that obtained by Minaev^37^ at the multireference CASSCF(16,11) level of theory. In Ref. ^38^, Silva computed the MECP for four different (1H)-3-hydroxy-4-oxoquinolines-O_2_ complexes, and found considerably different r_C2-O2_ depending on the substituent, ranging from 2.23 Å for QND to 1.56 Å when the methyl group is replaced by a fluoride. The energy of the MECP for the F substituted oxoquinoline was the smallest, 9.2 kcal/mol, 1 kcal/mol lower than in our calculations, also associated with a small r_C2-O2_ distance. Differences between our results and those in Ref. ^38^ can be associated to the different functional used (B3LYP vs M06-2X) or different simulation of the interactions with the solvent (PCM continuum model using water as a solvent vs QM/MM model). To check the effect of r_C2-O2_ on the energy of the MECP, we calculated the minimum of the crossing seam for different given r_C2-O2_ distances and we obtained that for r_C2-O2_ = 2.0 and 2.3 Å, the energies of the crossing point were only 1.8 kcal/mol and 2.7 kcal/mol above the MECP, respectively.

With respect to the SOC value, a value of 75 cm^-1^ was obtained for other similar spin-forbidden cofactor-less enzymatic reactions,^39^ leading to an increase of just 1.6 kcal/mol to the barrier, 1.5 kcal/mol lower than for the reaction between QND and O_2_. Assuming that this value is the upper-limit for a cofactor-less reaction involving O_2_, other points of the crossing seam (and not necessarily the MECP) could act as effective transition state if their energy is no more than 1.5 kcal/mol above the energy of the MECP, and show a SOC value closer to 75 cm^-1^. Even if that is the case, the effective barrier of the intersystem-crossing stage would be higher than for the CO release stage.

---

**Table S1.**
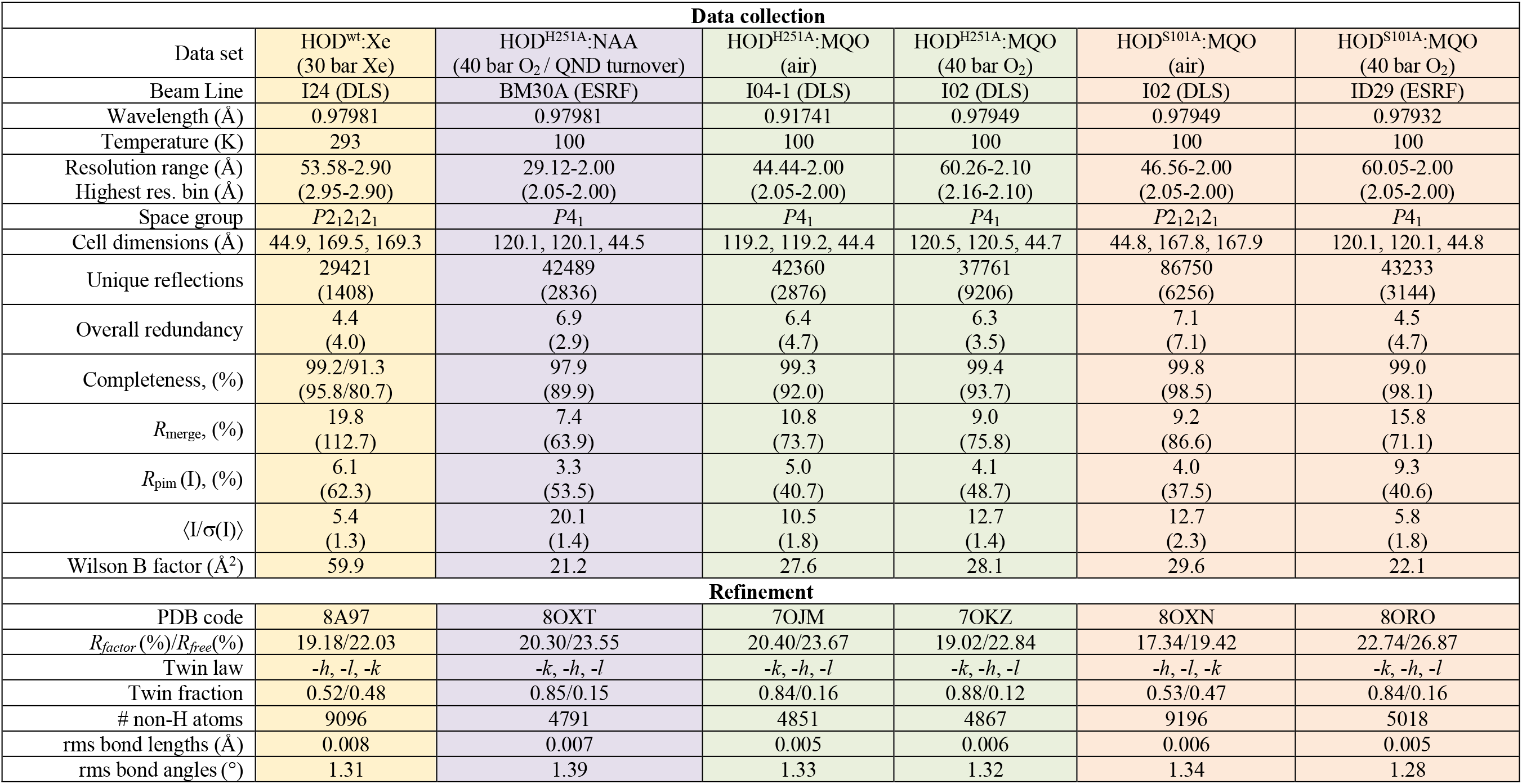
Data collection and refinement statistics.

**Figure S1.**
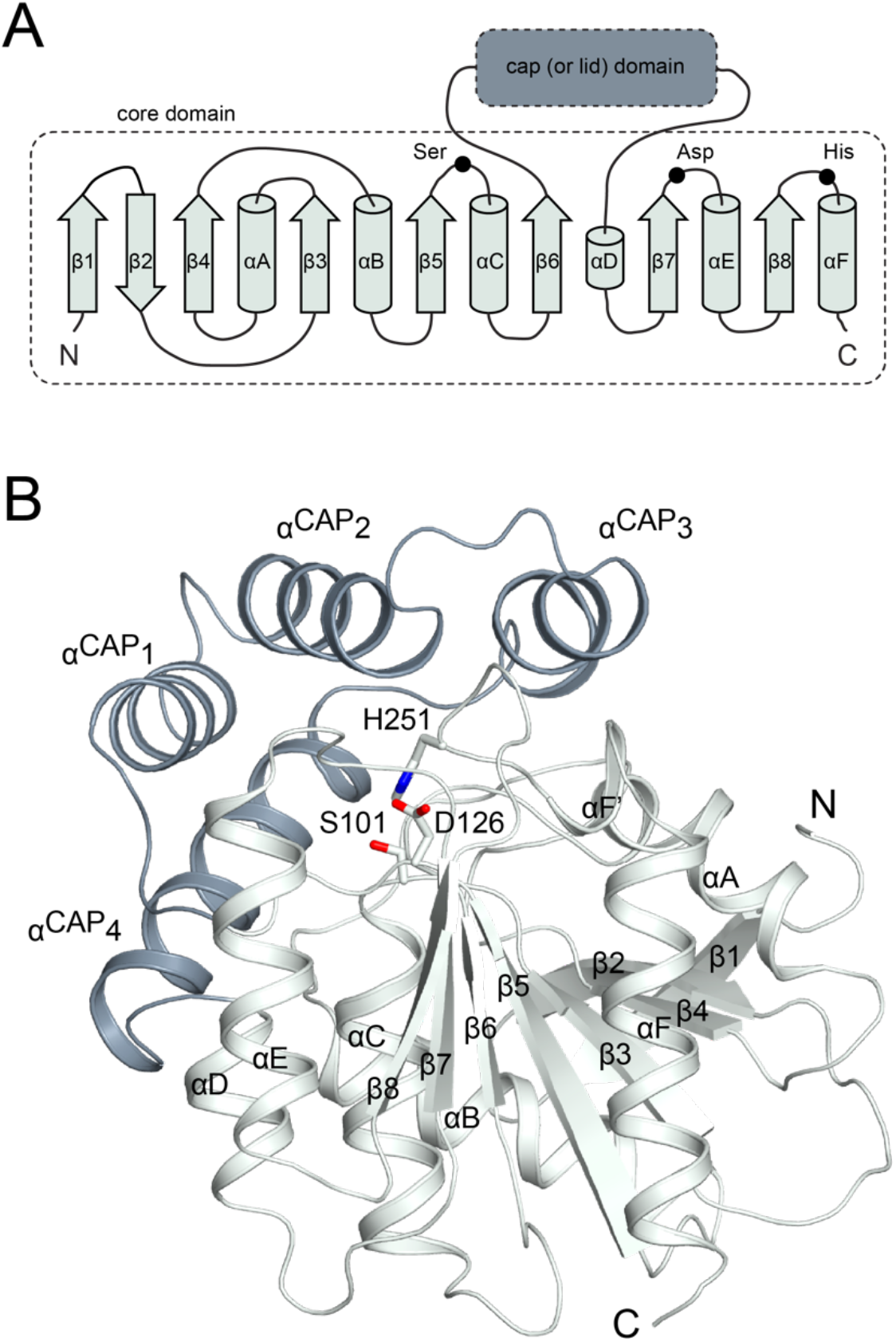
ABH fold. (A) General ABH fold topology and (B) ABH three-dimensional architecture exemplified by the crystal structure of *Arthrobacter nitroguajacolicus* Rü61a 1-*H*-3-hydroxy-4-oxoquinaldine 2,4-dioxygenase (HOD) (PDB code 2WJ3). HOD is the focus of the present study. The core domain of ABHs (here shown in light grey) is an eight-stranded, mostly parallel, β-sheet surrounded by α-helices. This is very often embellished by a cap or lid domain (here shown in dark grey) inserted between β6 and the short αD that provides structural and functional variation. In the case of HOD the cap domain is constituted by four α-helices. A very conserved feature of ABHs is the Ser (nucleophile), His, and Asp (acidic residue, sometimes replaced by Asn) catalytic triad represented by black dots in the topology diagram. S101, H251 and D126 form the catalytic triad in HOD. These residues are shown as sticks in panel B with nitrogen and oxygen atoms in blue and red, respectively. The nucleophile is hosted by the ‘nucleophilic elbow’, a tight turn connecting β5 and αC. In some ABH-fold dioxygenases the nucleophile is replaced by an alanine. The acidic residue, generally positioned after β7, is relocated after β6 in HOD. The catalytic histidine is near the C-terminus after the β8 strand. N and C indicate the protein N-terminus and C-terminus, respectively.

**Figure S2.**
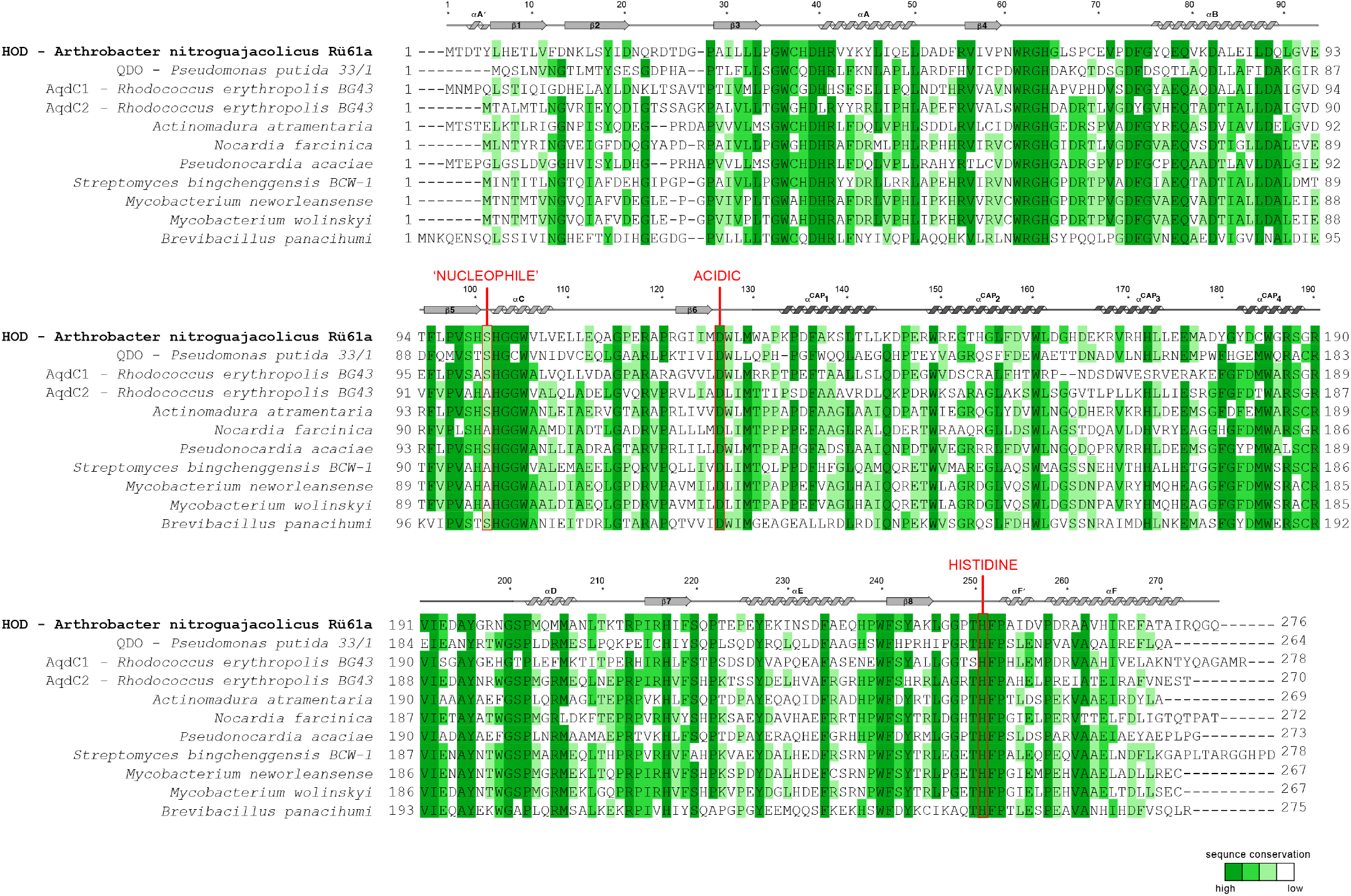
Sequence alignment of various predicted and validated bacterial ABH-fold cofactor-independent 2,4-dioxygenases exhibiting high similarity to *A. nitroguajacolicus* Rü61a 1-*H*-3-hydroxy-4-oxoquinaldine 2,4-dioxygenase (HOD). Residues are highlighted in different shades of green according to their sequence identity. Secondary structure elements for HOD are also shown for reference. Residues of the nucleophile-histidine-acidic catalytic triad are indicated. Whilst the His-Asp dyad is totally conserved, the nucleophile is not and, in several organisms, the latter residues is replaced by an alanine.

**Figure S3.**
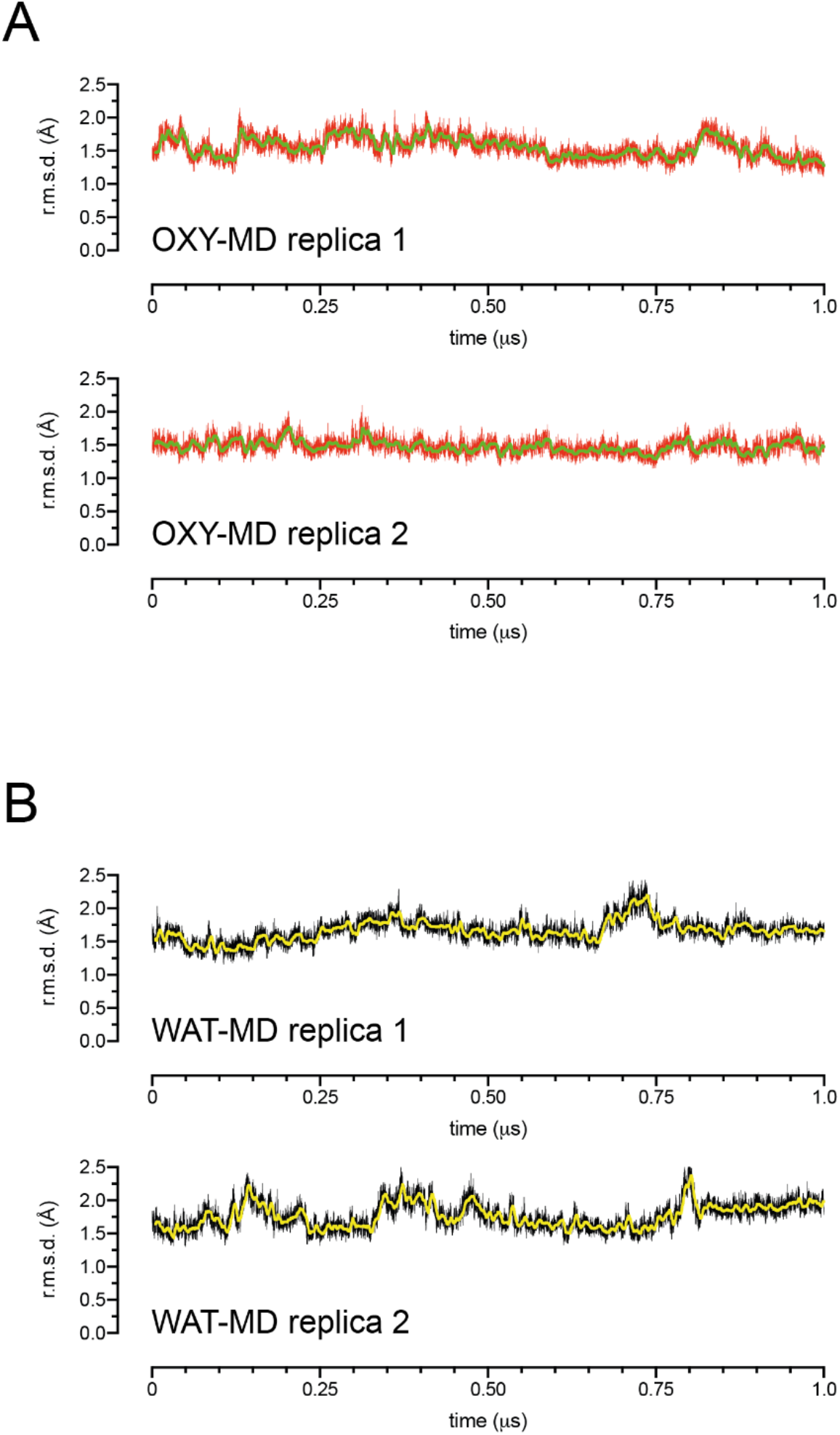
Root mean square deviation (r.m.s.d.) from the starting equilibrated crystallographic structure for HOD during 1 μs-long replicas Molecular Dynamics (MD) simulations. OXY-MD runs were carried out in the presence of ten O_2_ molecules in a solvated box of approximately 74×74×74 Å^3^. WAT-MD did not include O_2_ molecules in the simulations. Sampling was performed every 50 ps of each run. Thick lines represent the moving average calculated using a 5 ns window.

**Figure S4.**
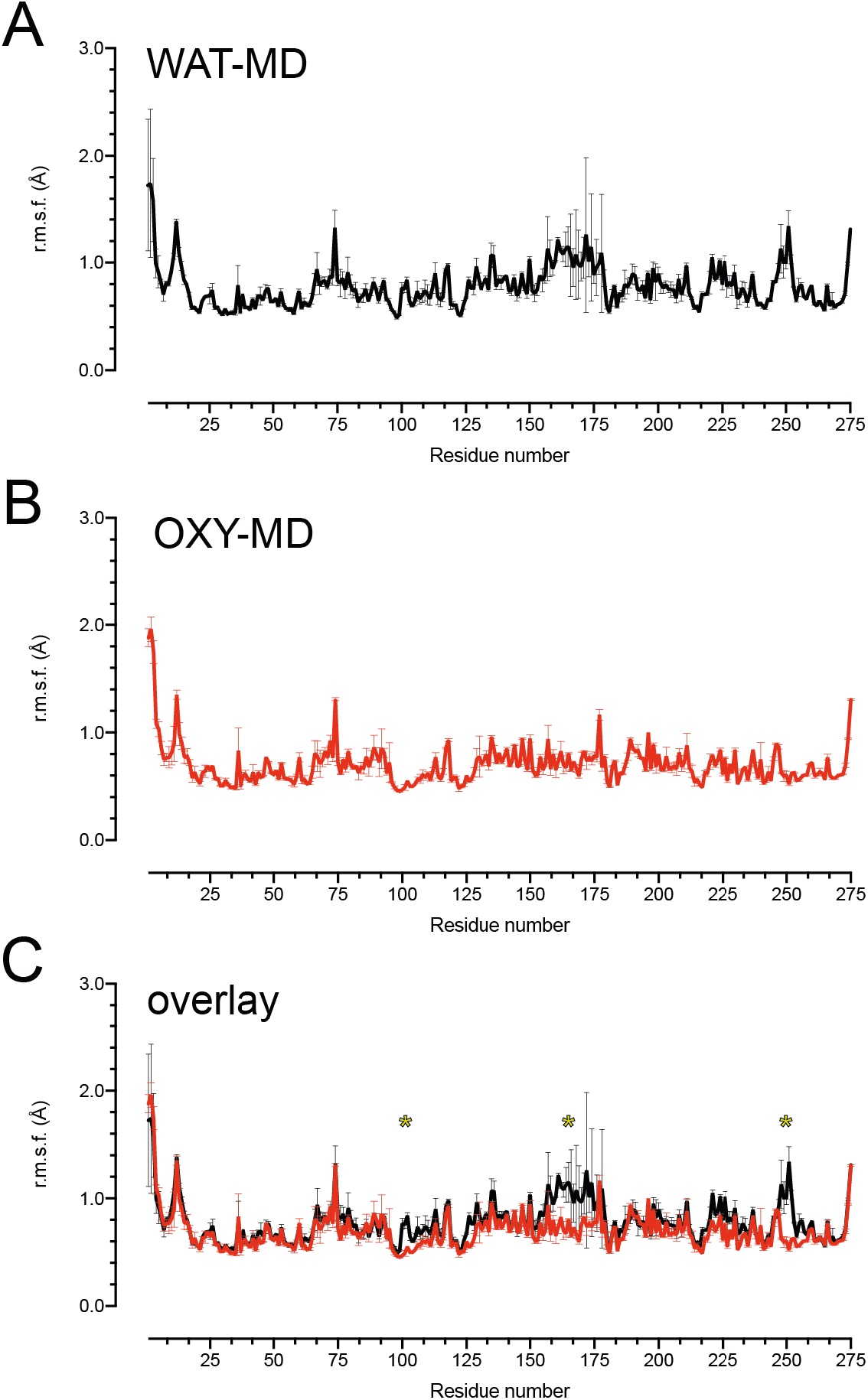
Root mean square fluctuations (r.m.s.f.) from the average structure for HOD during WAT-MD (A) and OXY-MD (B) simulations. OXY-MD runs were carried out in the presence of ten O_2_ molecules to a solvated box of approximately 74×74×74 Å^3^. WAT-MD did not include O_2_ molecules in the simulations. Two replica 1-μs simulations were run for each system and data are presented as average r.m.s.f. with standard deviations. (C) is an overlay of A and B with asterisks highlighting the regions with different mobility.

**Figure S5.**
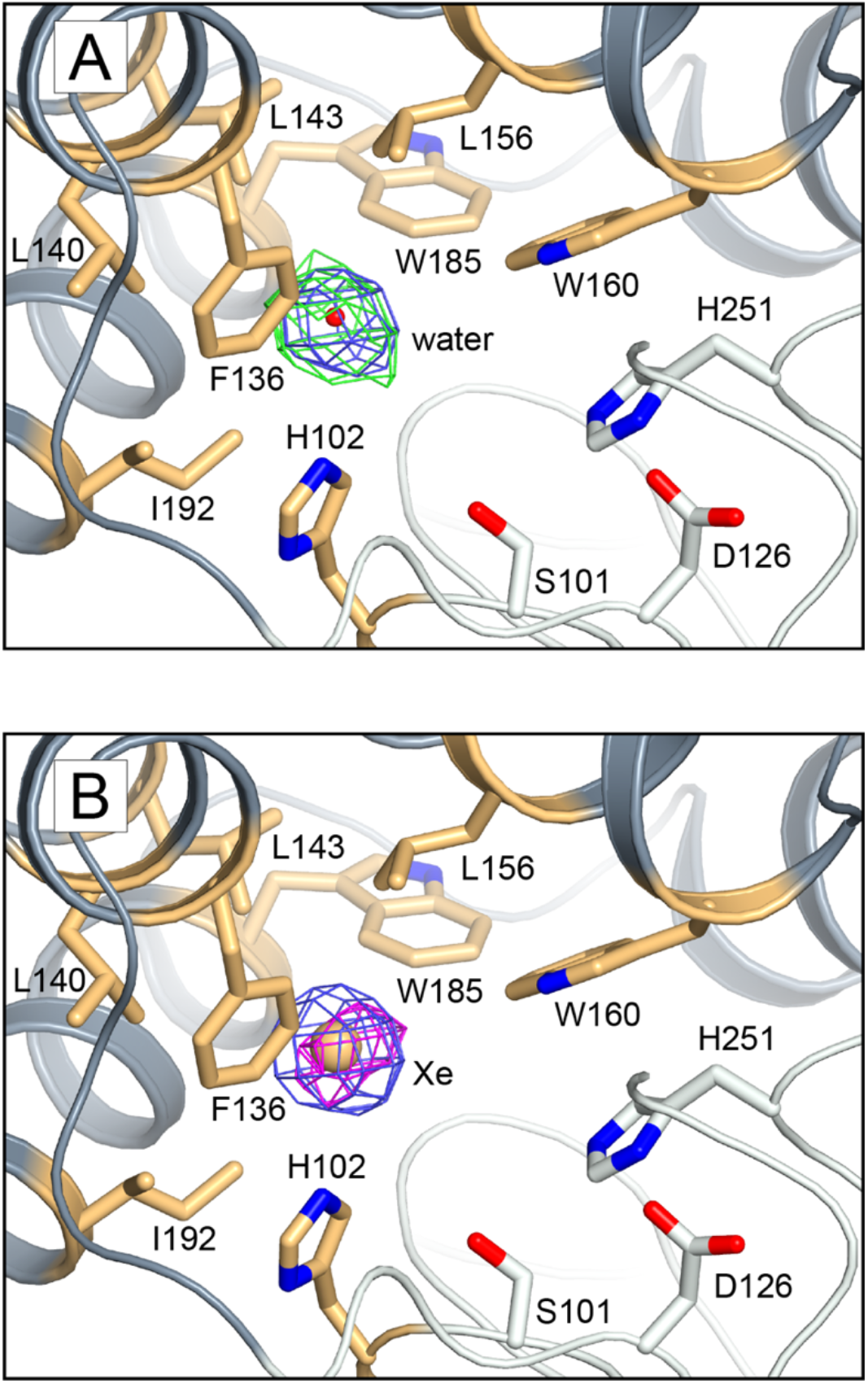
Validation of Xe binding at the *B*-site. (A) Interpretation of the Xe site as a water molecule results in significant residual density in Fourier difference maps. Post-refinement 2*mF*_o_-*DF*_c_ and *mF*_o_-*DF*_c_ electron density maps are shown at the +1.0σ (blue) and +3.0σ (green) levels, respectively. (B) Interpretation as a Xe is satisfactory and occupancies refine between 0.4 and 0.6 in the four different molecules in the a.u. The 2*mF*_o_-*DF*_c_ electron density map is shown at the +1.0σ level in blue. The map displayed in magenta is the NCS-averaged anomalous difference map contoured at +5.5σ. We note that the anomalous signal for this experiment is weak as the wavelength used for data collection was far from the f” peak to minimize radiation damage at room temperature. Also, data at 2.9 Å-resolution exhibit non-merohedral twinning (twinning fractions 0.52/0.48) which negatively affects the anomalous signal. Despite these limitations the interpretation of this site as Xe is unambiguous. Hydrophobic residues stabilizing the Xe atom are highlighted in gold and shown as sticks. The catalytic triad (S101/H251/D126) is also shown.

**Figure S6.**
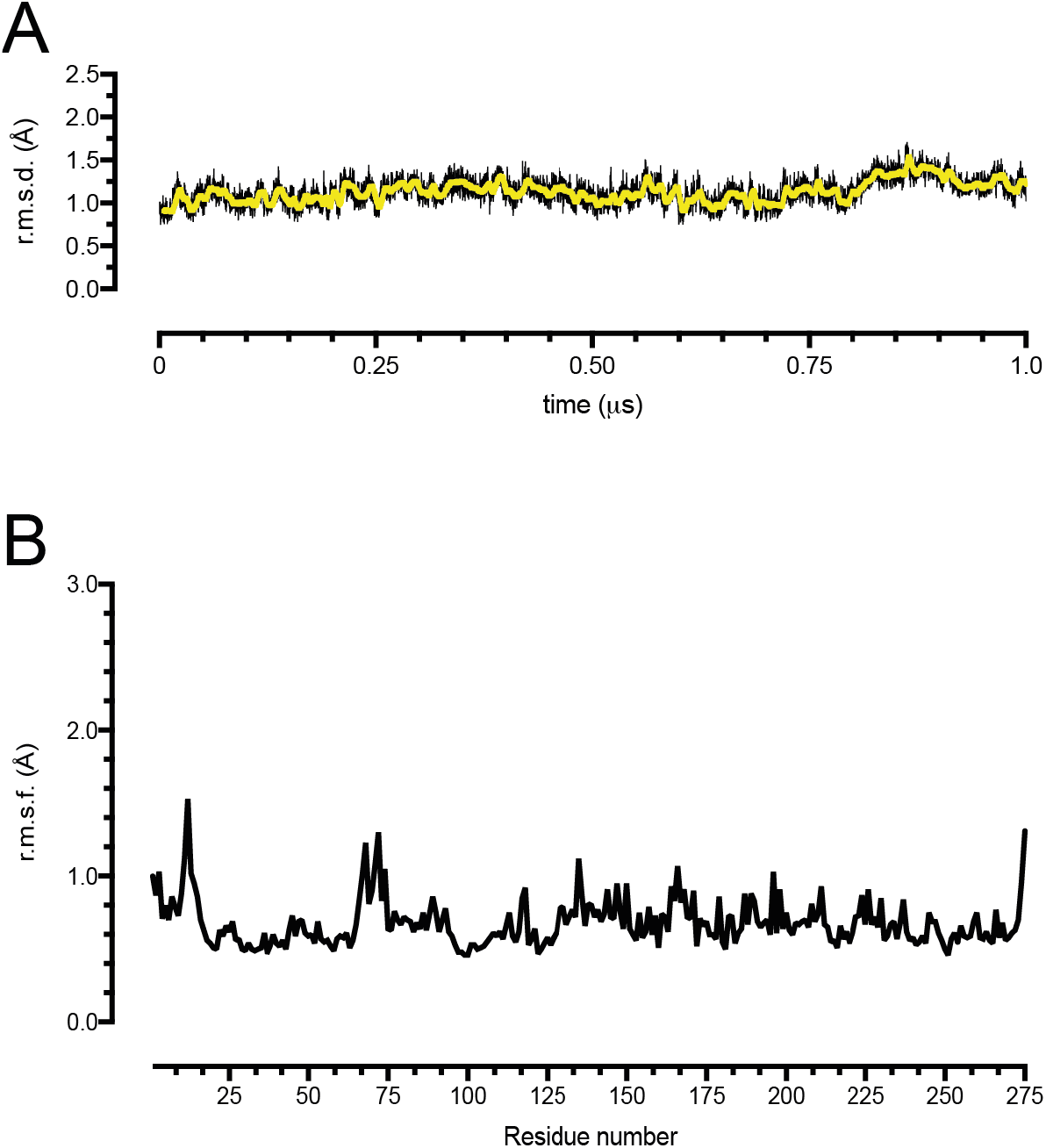
Molecular Dynamics (MD) simulation of the HOD-QND complex. (A) Root mean square deviation (r.m.s.d.) from the starting equilibrated crystallographic structure for the HOD-QND complex (PDB code 2WJ4), during a 1 μs-long MD simulation. Sampling was performed every 50 ps of each run. The thick yellow line lines represent the moving average calculated using a 5 ns window; (B) root mean square fluctuation (r.m.s.f.) from the average structure during the simulation.

**Figure S7.**
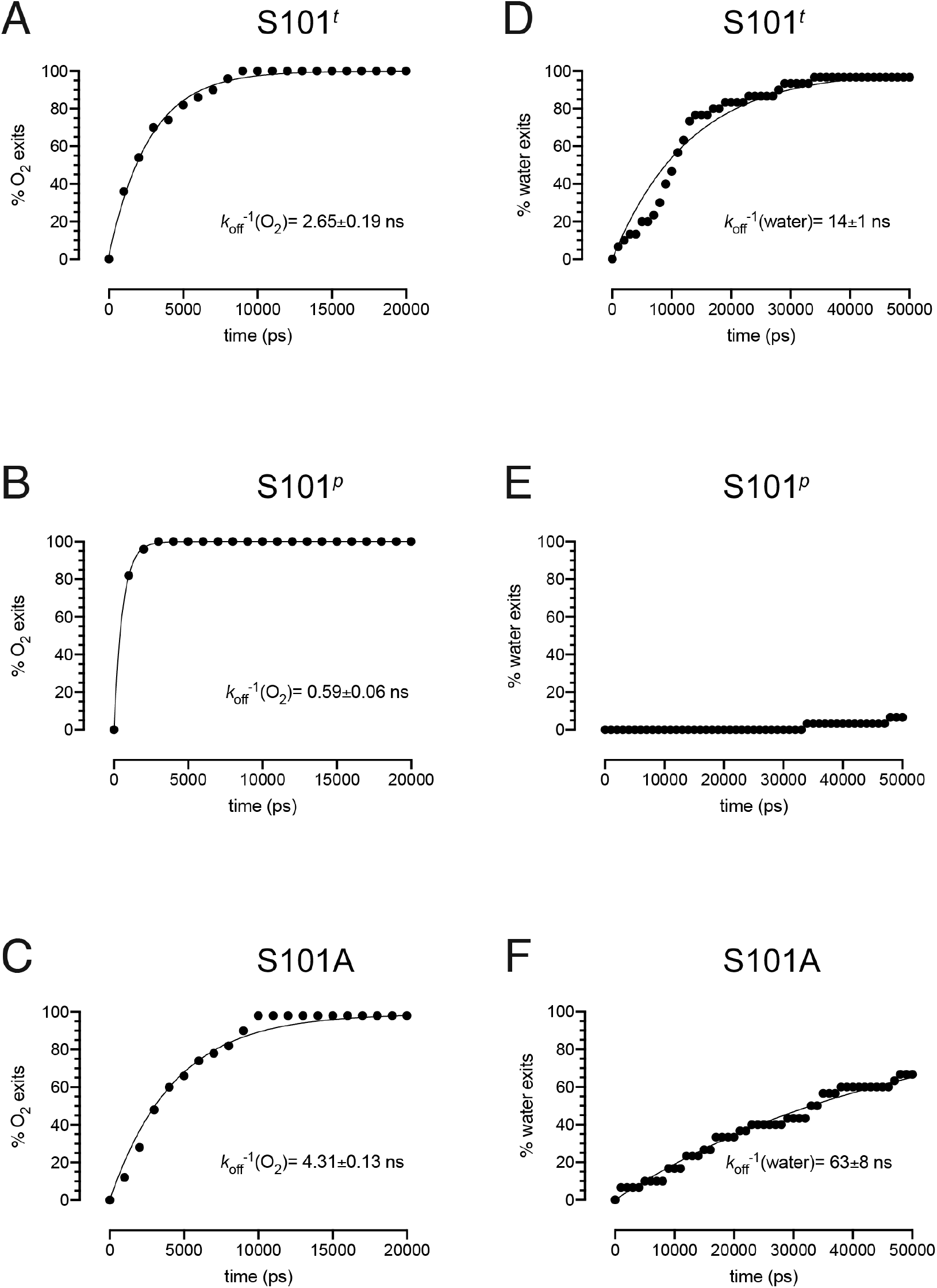
Fitting of residence times for O_2_ (A-C) or water (D-F) at the *R*-site with S101 restrained to either the *trans* rotamer (A,D), or to the *plus* rotamer (B,E), or, alternatively, with S101 is replaced by an alanine (C,F). The cumulative distributions were fitted to the equation (2) to determine the residence times (*λ* = *k*_off_^-1^). In the case of (E) fitting is not possible as there are too few escape events with 50 ns.

**Figure S8.**
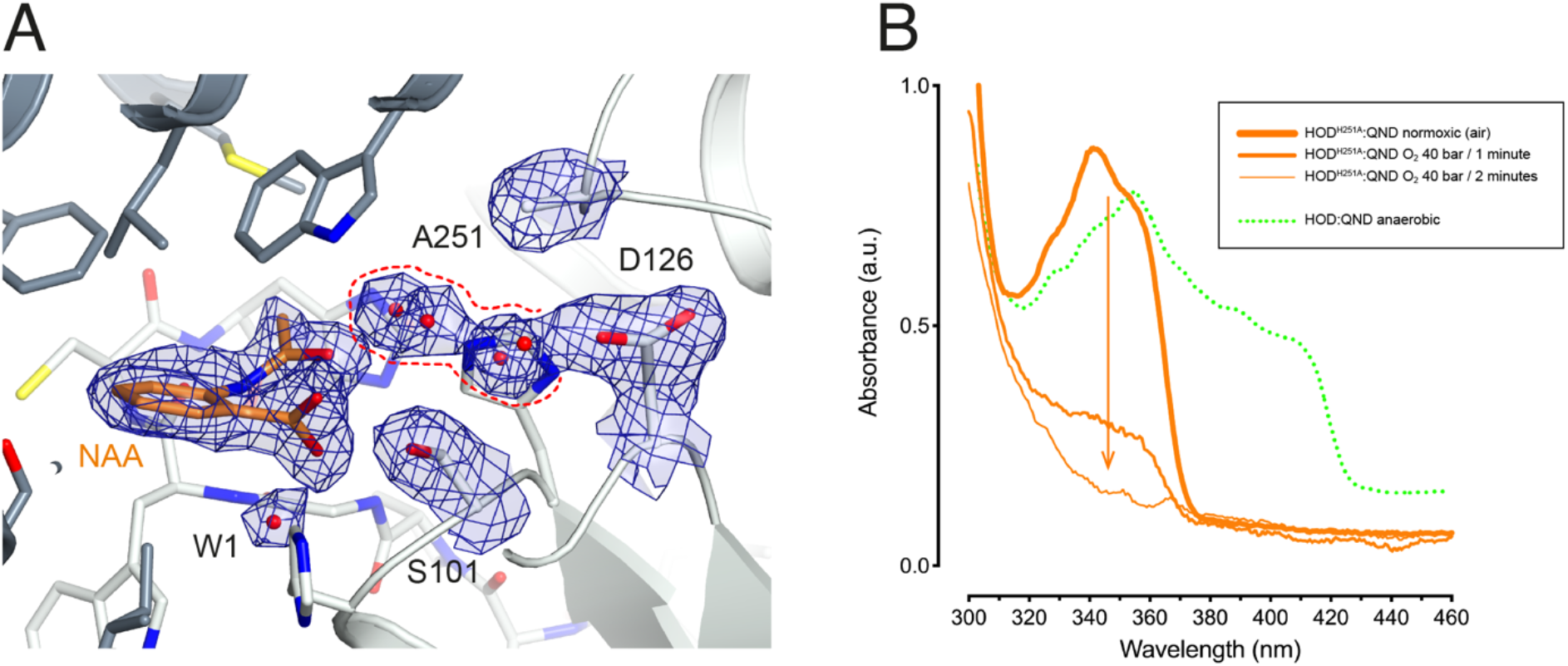
*In crystallo* turnover. (A) Active site of the HOD^H251A^-QND complex following O_2_ pressurization at 40 bar for 2 minutes. QND is fully converted to *N*-acetylanthranilate (NAA), the product of the HOD-catalyzed reaction. Electron density maps (2*mF*_o_-*DF*_c_) are shown at the +1.0σ level for NAA, the side chains of S101, D126, A251 (catalytic triad), the water molecule at the *R*-site (W1) and the solvent network (highlighted by the red dotted line) that connects D126 to the organic ligand. *in crystallo* UV-visible spectra of HOD^H251A^-QND under normoxic conditions (thick orange line), after 1-minute O_2_ pressurization (medium thickness orange line), after 2-minutes O_2_ pressurization (thon orange line). The *in crystallo* UV-visible spectrum of the anaerobic HOD-QND complex (dotted green line) is also shown for reference. A small offset was added to the green trace to improve clarity. Pressurization enables O_2_ access to the *R*-site in the pre-formed HOD^H251A^-QND complex promoting turnover without buildup of the substrate anion. The structure suggests that the marginal residual catalytic activity of the HOD^H251A^ variant (*k*_cat_ = 0.0006 s^-1^ at pH 6.5 compared to 16.3 s^-1^ for wtHOD) relies on solvent-mediated substrate deprotonation.

**Figure S9.**
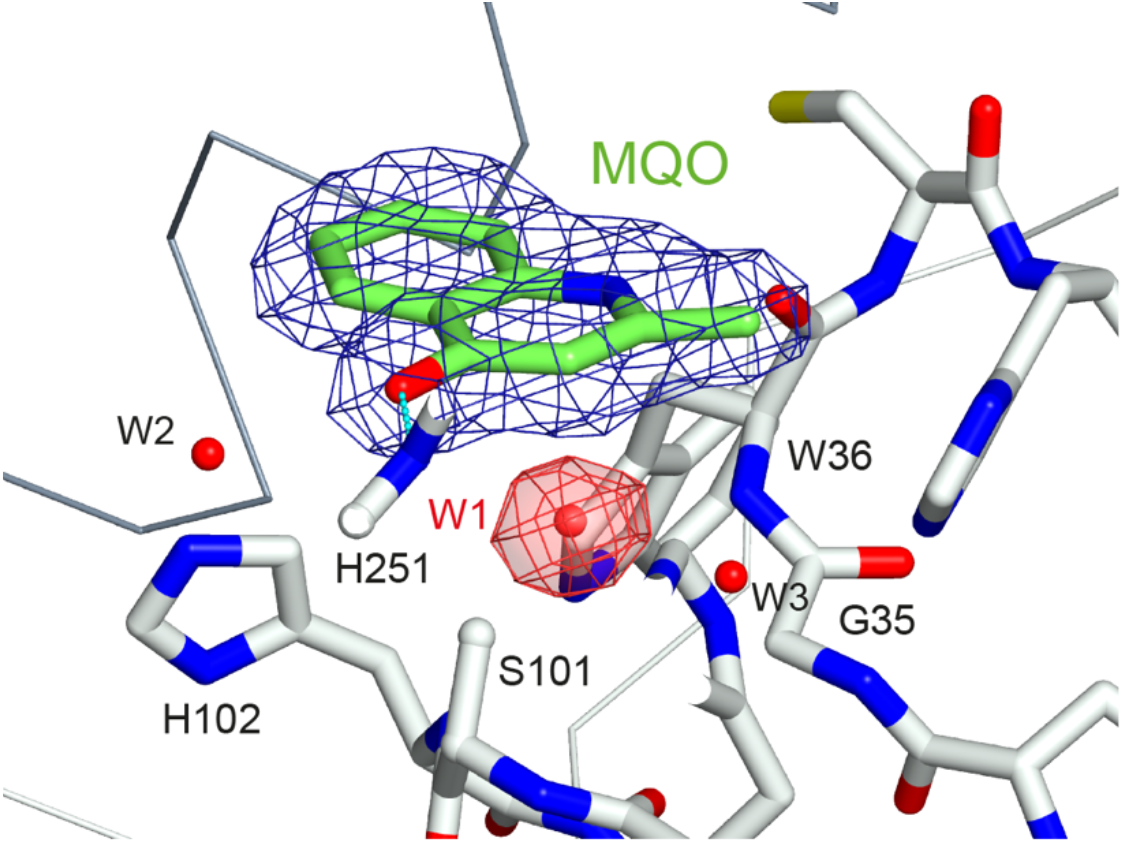
Representative 2*mF*_o_-*DF*_c_ electron density for the HOD^S101A^-MQO complex under normoxic conditions. Electron density maps are shown at the +1.0σ level as chicken-wire representation for MQO (blue) and a water molecule (W1) at the *R*-site. The latter is displayed in red for clarity. The view is the same as in Figure 5B of the main text.

**Figure S10.**
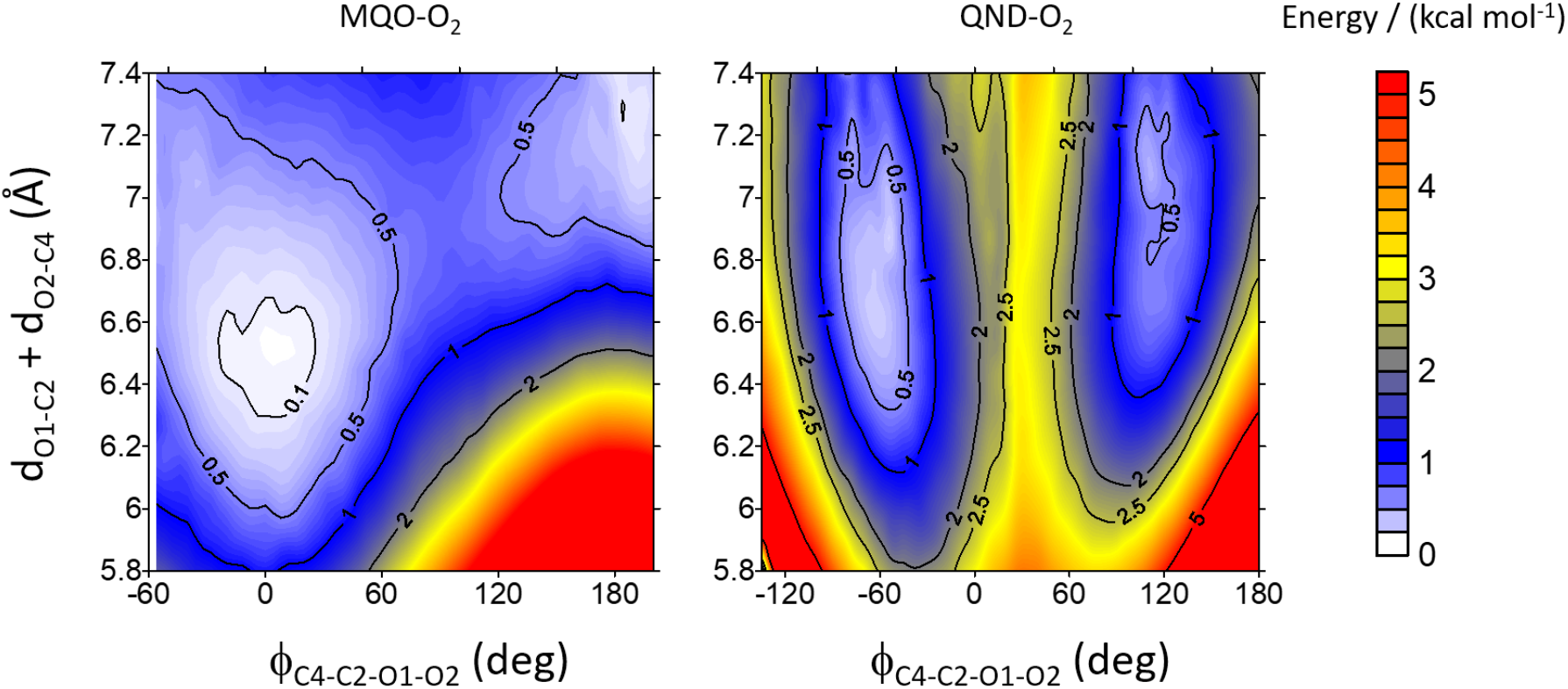
3D contour maps for the energy profiles for the O_2_-ligand (MQO on the left and QND on the right) complexes as a function of the dihedral angle Φ = (C4-C2-O2-O1), and the sum of (C2-O1) and (C4-O2) distances. The latter was used as a restraint in the calculations for the isolated complexes. Around the equilibrium geometries, the energy is relatively insensitive to small changes in the restraint value. The two minima for the MQO-O_2_ complex at Φ=0 and 180° represents the same structure, reflecting that O1 and O2 are equivalent.

**Figure S11.**
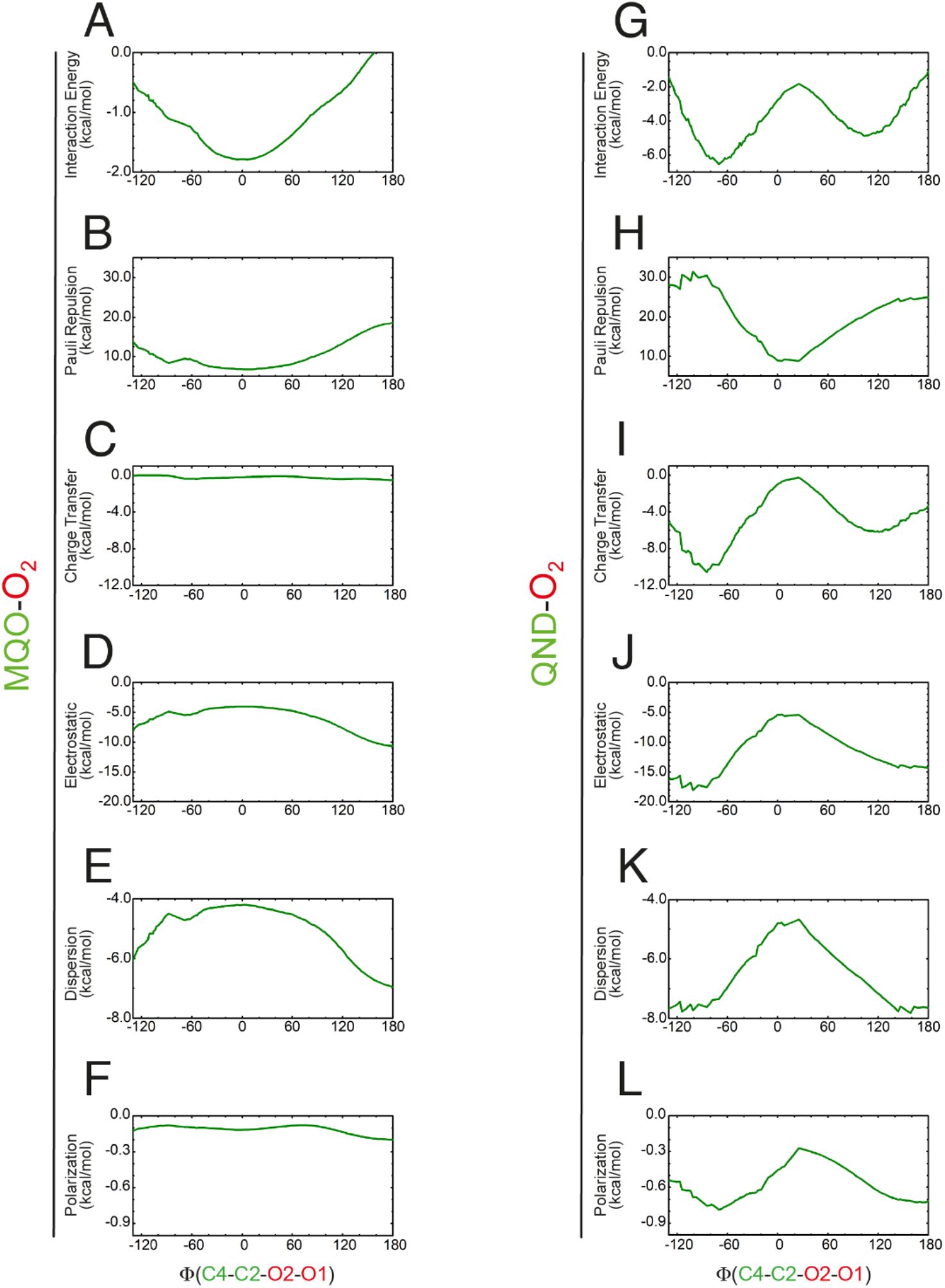
Energy decomposition analysis for the MQO-O_2_ (A-F) and QND-O_2_ (G-L) complexes as a function of the dihedral angle Φ=(C4-C2-O2-O1). Interaction energy values (A,B) are the sum of their respective Pauli repulsion (B,H), charge transfer (C,I), electrostatic (D,J), dispersion (E,K), and polarization (F,L) terms (A=B+C+D+E+F, G=H+I+J+K+L).

**Figure S12.**
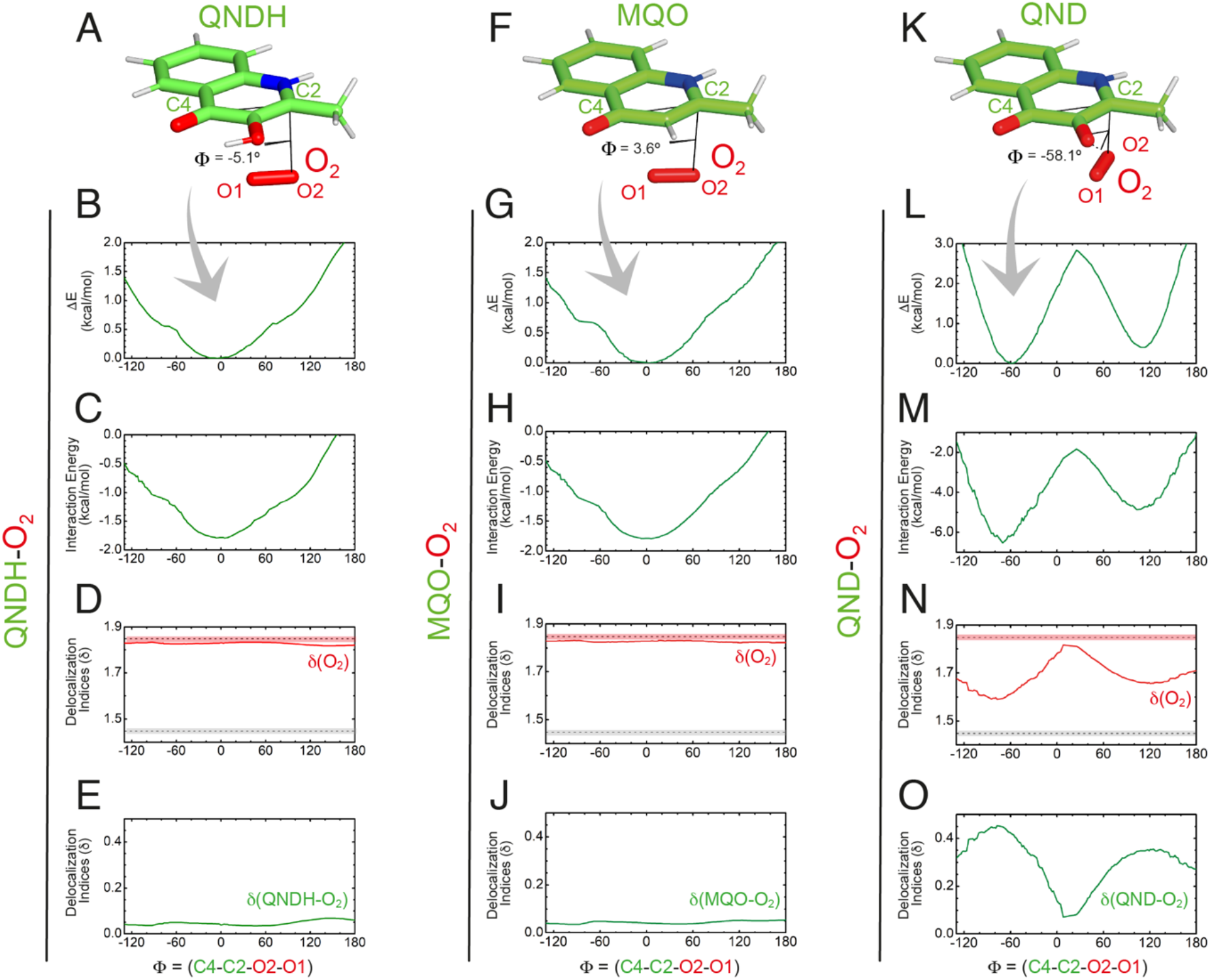
QM geometry restrained optimization and delocalization indices for QNDH-O_2_ (A-E), MQO-O_2_ (F-J) and QND-O_2_ (K-O) complexes as a function of the dihedral angle Φ = (C4-C2-O2-O1). Panels (F-O) are identical to panels (C-L) in Figure 5 of the main text and are shown here for reference. In panels (D,I,N) delocalization indices for isolated O_2_ (dashed red line) and the aromatic C-C bond in benzene (dashed grey line) are given for reference. Protonation of QND (QNDH) results in an overall neutral system QNDH-O_2_ system that behaves like MQO-O_2_.

**Figure S13.**
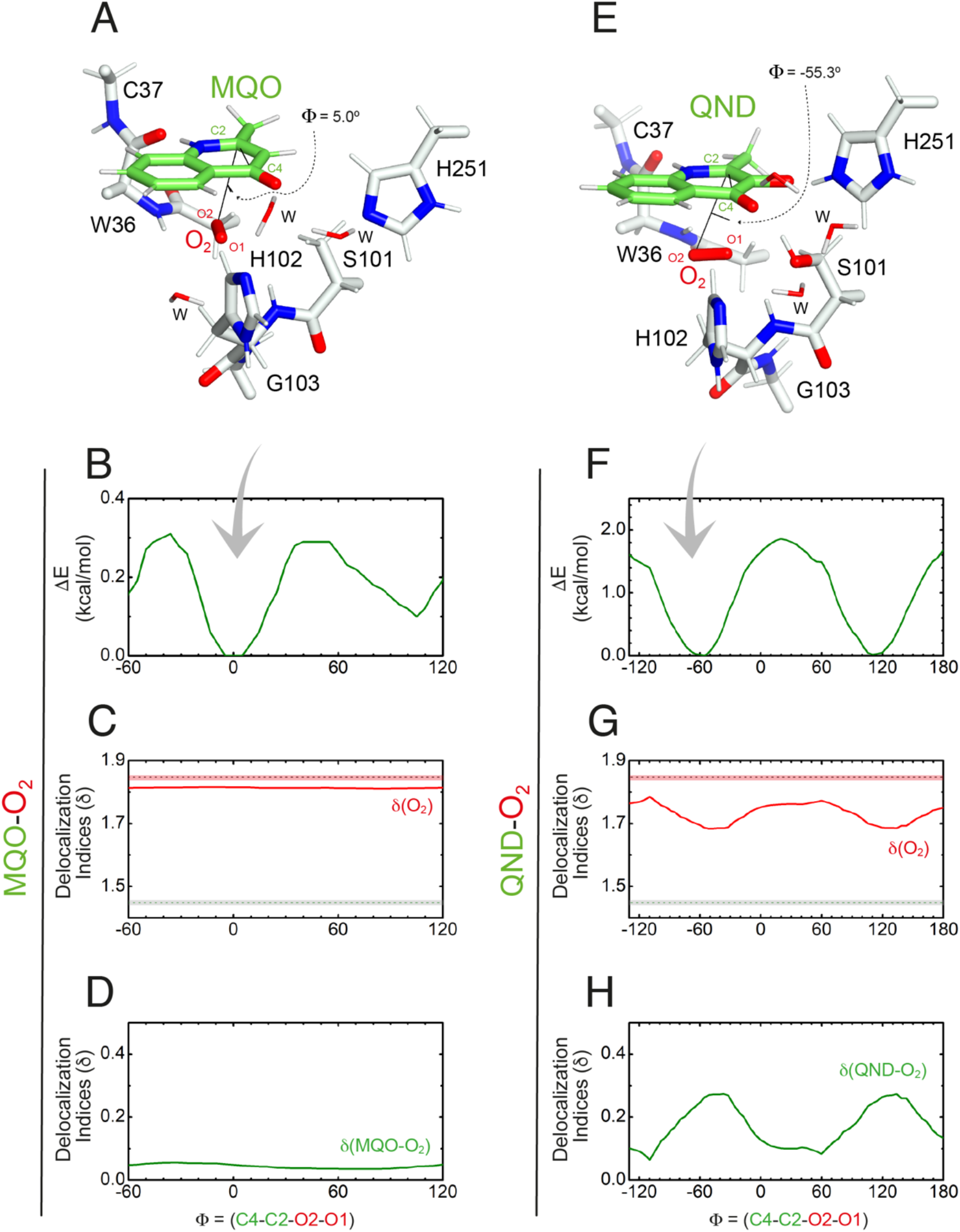
QM/MM geometry restrained optimization and delocalization indices for the MQO-O_2_ (A-D) and QND-O_2_ (E-H) complexes as a function of the dihedral angle Φ = (C4-C2-O2-O1). (A,E) are stick representations of the minimized structures. (B,F) show the total energy of the complexes as a function of Φ. (C,G) display the delocalization index (δ) for the O-O bond in the complex. For comparison, the dashed red and grey lines represent the delocalization indices for O-O and C-C bonds in isolated O_2_ and in benzene, respectively. (D,H) show the delocalization index for the interaction between the substrate and O_2_.

**Figure S14.**
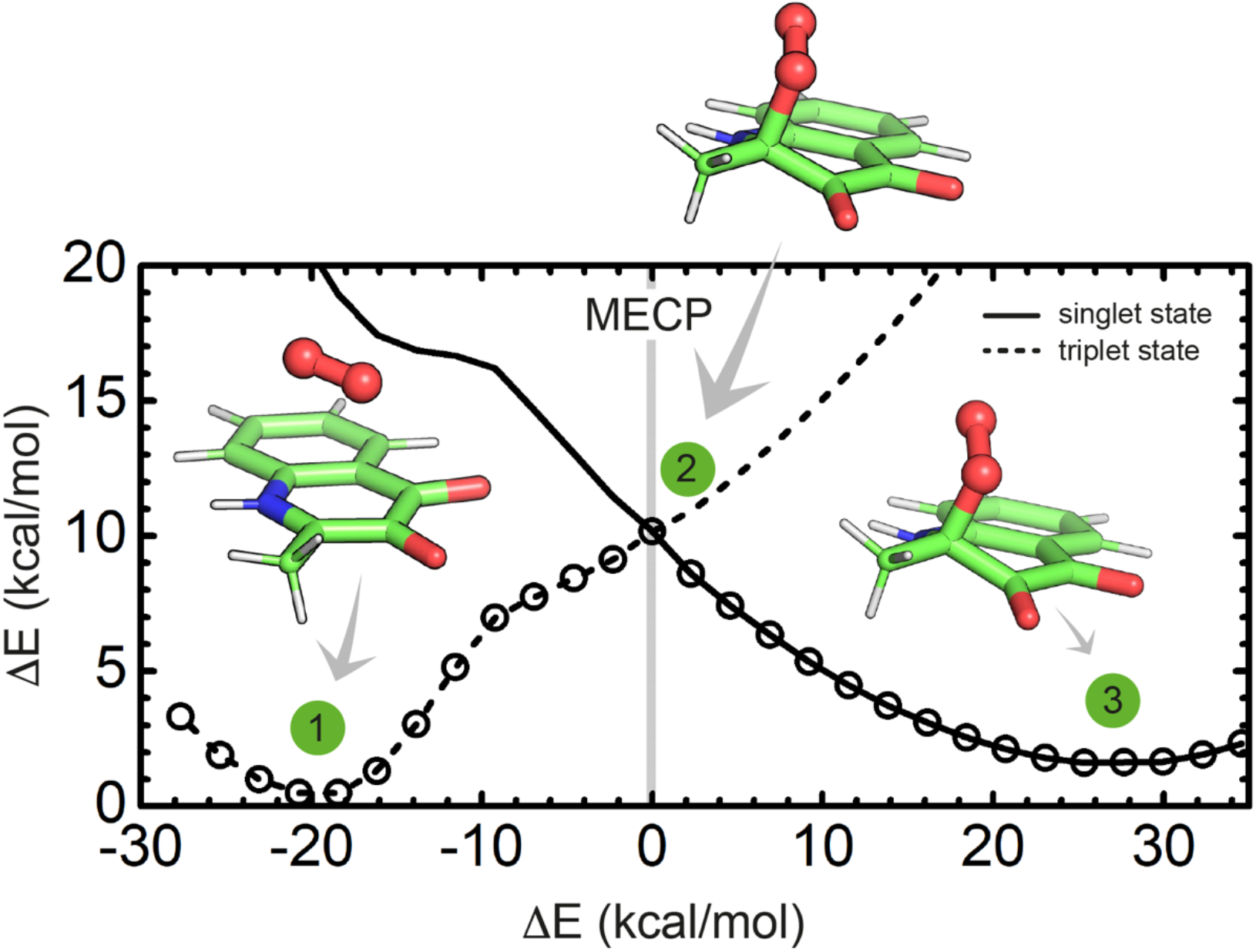
Energy profile for the transition from the triplet to the singlet state and formation of the peroxide intermediate. The energy is presented as a function of the energy difference between the triplet and the singlet state, which was chosen as a reaction coordinate. The relevant structures at the points indicated are shown as stick (ball-and-stick for O_2_) representations. Structure 2 represents the minimum energy crossing point (MECP).

**Figure S15.**
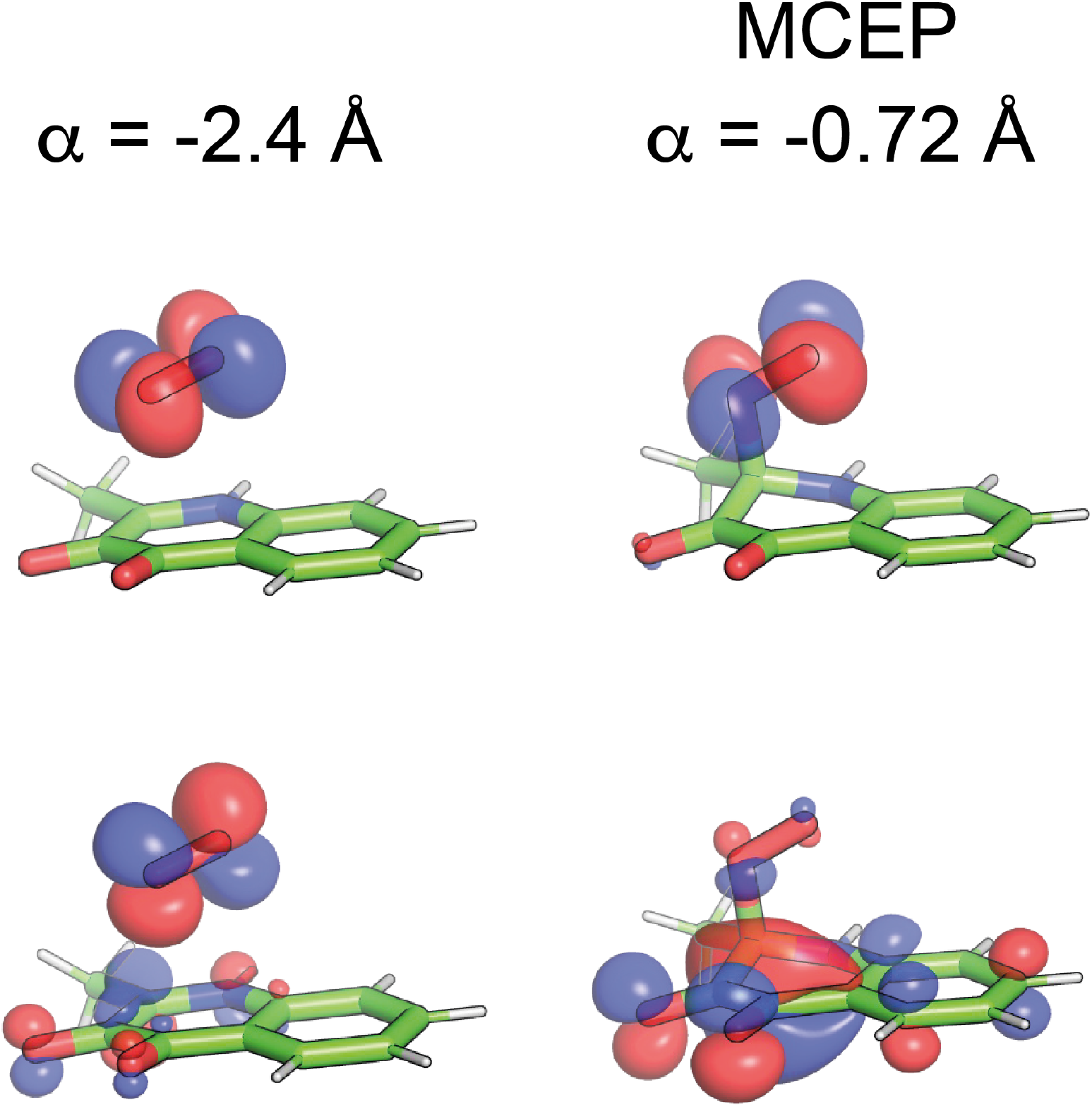
Shape of the half-occupied orbitals for the QND-O_2_ complex at the reactants asymptote (left panels), and at the MECP (right panels). The orbitals were localized following the procedure described in Ref. ^40^ At the reactants asymptote, half-occupied orbitals correspond to the two π* orbitals of O_2_. The π* perpendicular to the π cloud of QND (π_x_*), is not mixed with any QND orbital. However, the π* orbital that stacks with the π cloud of QND (π_y_*) is significantly mixed with the π cloud of QND. As QND and O_2_ approach each other, mixing is more substantial, leading to an occupied orbital with bonding character along the C2-O bond, and a higher energy orbital that shows an antibonding character both along O1-O2 and C2-O bonds. The latter orbital is singly occupied in the triplet state but it is a virtual orbital in the singlet state, explaining why the singlet state is stabilized along the reaction path while triplet state is destabilized

